# An improved organ explant culture method reveals stem cell lineage dynamics in the adult *Drosophila* intestine

**DOI:** 10.1101/2021.12.17.473114

**Authors:** Marco Marchetti, Chenge Zhang, Bruce A. Edgar

## Abstract

In recent years, live-imaging techniques have been developed for the adult midgut of *Drosophila melanogaster* that allow temporal characterization of key processes involved in stem cell and tissue homeostasis. However, current organ culture techniques are limited to imaging sessions of ≤16 hours, an interval too short to track dynamic processes such as damage responses and regeneration, which can unfold over several days. Therefore, we developed a new organ explant culture protocol capable of sustaining midguts *ex vivo* for up to 3 days. This was made possible by the formulation of a culture medium specifically designed for adult *Drosophila* tissues with an increased Na^+^/K^+^ ratio and trehalose concentration, and by placing midguts at an air-liquid interface for enhanced oxygenation. We show that midgut progenitor cells can respond to gut epithelium damage *ex vivo*, proliferating and differentiating to replace lost cells, but are quiescent in healthy intestines. Using *ex vivo* gene induction to promote stem cell proliferation, we demonstrate that intestinal stem lineages can be traced through multiple cell divisions using live imaging. Both asymmetric and symmetric divisions can be identified in the reconstructed lineages. We find that daughter cells of asymmetric divisions remain in close proximity of each other, while the progeny of symmetric divisions actively move apart, with implications for cell differentiation and tissue organization. We show that the same culture set-up is useful for imaging adult renal tubules and ovaries for up to 72 hours. By enabling both long-term imaging and real-time *ex vivo* gene manipulation, our simple culture protocol provides a powerful tool for studies of epithelial biology and cell lineage behavior.

## INTRODUCTION

Endo-and ectodermal epithelia comprise essential interfaces between an organism and its environment. As such, they form a first line of defense that is frequently subjected to diverse types of insult. This situation requires that epithelia be able to mount appropriate responses. This is possible in part due to the action of resident stem cells which, through their ability to self-renew and produce differentiated progeny, allow epithelia to regenerate both structurally and functionally. The adult *Drosophila melanogaster* midgut is a prime example of this as its population of intestinal stem cells (ISCs) are able to interpret signals from their surrounding environment such as cytokines released by neighboring damaged enterocytes (EC)^1–5^. When this interaction occurs, normally quiescent stem cells rapidly respond to the needs of the tissue, proliferating and stimulating the differentiation of their progeny to replace lost cells, thus repairing the damaged epithelium^3,5^.

The understanding of epithelial biology has been greatly advanced by protocols for the *ex vivo* culture and imaging of tissues and organs. For example, mammalian intestinal organoids have greatly advanced the field by easily allowing the direct observation of stem cell behavior, without the need for intravital imaging^6^. Several protocols have been developed for the live-imaging of *Drosophila* tissues and organs such as imaginal discs^7–11^, larval brains^12,13^, ovaries^14–16^, testis^17^ and, more recently, adult midguts^18–22^. The small size of fruit flies makes it possible to culture whole intact organs.

However, in contrast to mammalian tissues, many of which are easily cultured for long periods, most *Drosophila* organ cultures are limited in time to less than a day. This reflects an incomplete understanding of the culture conditions required to fully sustain explanted *Drosophila* tissues. Current approaches for the live-imaging of the fly midgut are limited to 16h of imaging due to the poor survival of explanted tissues^21,22^. Moreover, temperature-sensitive gene expression, knock-down, and knock outs, some of *Drosophila* genetics strongest assets, cannot currently be implemented in combination with extended live-imaging because the elevated temperature further limits tissue viability^21^.

To address these limitations, we developed an improved *ex vivo* culture system for the live-imaging of adult *Drosophila* midguts. Our culture system employs a novel tissue culture medium tailored to the needs of adult *Drosophila* cells and organs, and culture at an air-media interface to ensure optimal oxygenation. The technique has a straightforward design, allowing multiple samples to be prepared quickly and reproducibly. The setup allows the researcher to image up to 12 midguts simultaneously during live-imaging sessions of 48-72h. As the guts are fully explanted from the animal, every region of the organ is clearly available for imaging, thus expanding the number of questions that can be addressed. We show that, while in healthy explanted intestines progenitor cells are quiescent, midguts can still respond to damage ex vivo, with progenitors proliferating and differentiating in response to tissue damage Our protocol can also be used in conjunction with temperature-sensitive gene expression or knock-down. We demonstrate this by genetically driving stem cell proliferation. Moreover, due to the extended live-imaging window our protocol allows, we were able to follow cells undergoing multiple rounds of mitosis. In the reconstructed lineages we observed both symmetric and asymmetric mitotic events and found the behavior of their progeny to be distinct. By combining a long 48-72h imaging window and the possibility to use advanced *Drosophila* genetic tools, we provide a useful tool to probe and understand the biology of epithelial tissues.

## RESULTS

### A system for the long-term culture of adult *Drosophila* midguts

Current live-imaging protocols for the adult *Drosophila* midgut are limited to 16h imaging sessions^18–22^. To extend the survival of midguts *ex vivo* we developed a novel culture setup. The method is based on common available techniques for the culture of adult organs^6,9,16,22^ but it includes a refinement o several steps: 1) the dissection procedure was optimized to reduce tissue damage; 2) explanted organs are cultured in a sandwiched structure of agarose, rather than in a dome; 3) midguts are placed at a liquid-air interface for improved oxygenation; 4) the culture media has been adjusted to better approximate adult hemolymph. Please see the Methods section step-by-step descriptions of the procedure.

We found that the dissection technique is a key parameter in extending the viability of explanted midguts. Indeed, any stress (*e*.*g*. pulling and thus stretching the gut, pinching, etc…) introduced during dissection results in structural damage which lead to breaks of the epithelium during prolonged culture. Hence, we have optimized the dissection procedure to limit the handling of the midgut and thus the risk of damaging it (Figure 1A-N and Video 1).

**Figure 1:**
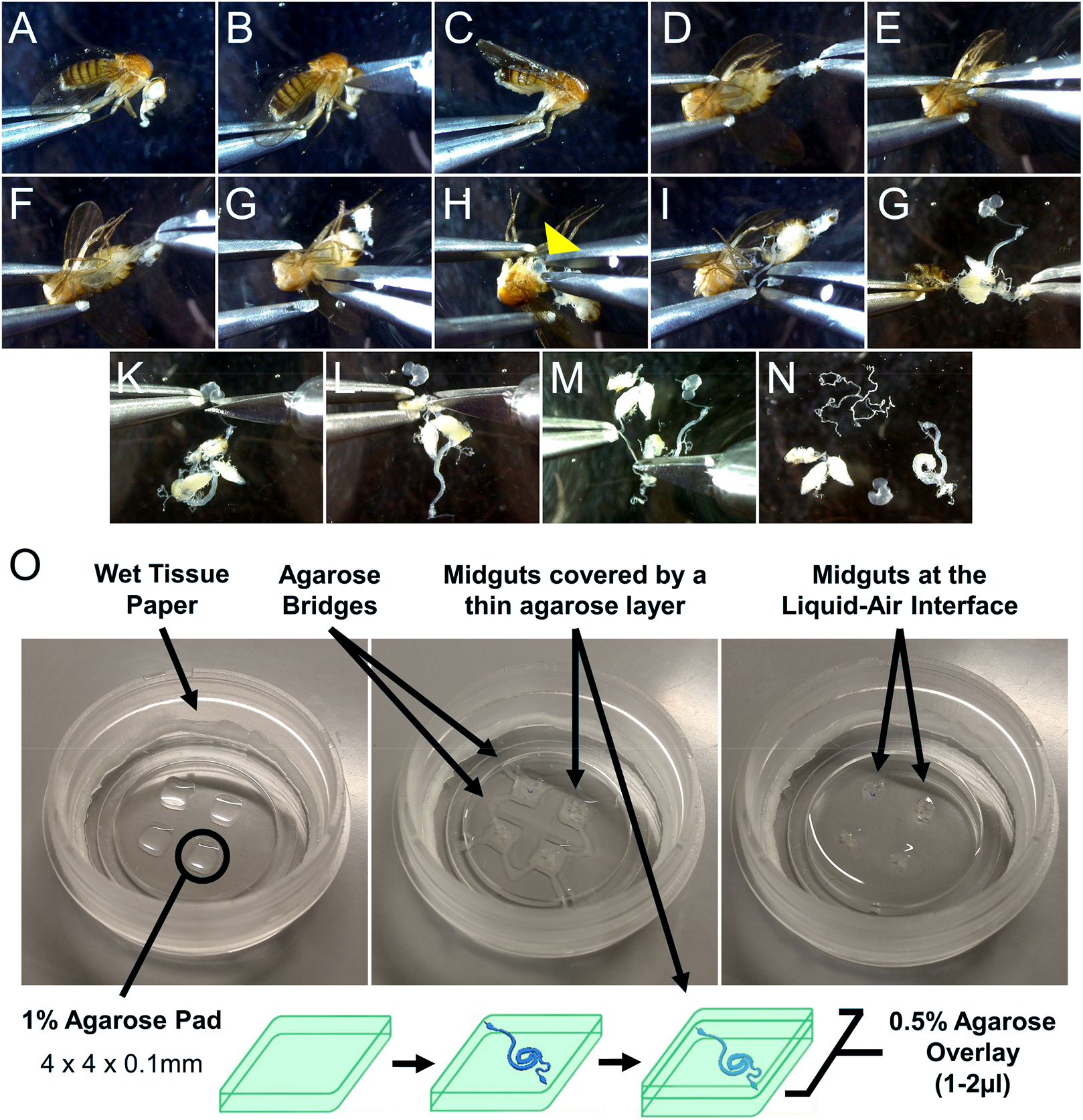
Sample preparation for live-imaging. (A-N) Minimal handling dissection used to gently explant adult Drosophila midguts, limiting the risk of damaging the intestines. See also Video 1. (O) Culture chamber setup (left) and mounting of explanted midguts (middle and bottom diagram) to produce an air-liquid interface culture (right). See methods section and Video 1 for in-depth description.

A key element of our system is enveloping explanted organs in a sandwich of low-melt agarose (Figure 1O). Immobilization in gels is a common solution for *ex vivo* culture of organs and tissues, and is especially important to provide stability for live-imaging applications^6,9,16,22^. For the adult *Drosophila* midgut specifically, an agarose gel also minimizes the effects of peristaltic movements, which, if uncontrolled, will impair imaging and can lead to epithelial tearing. Our approach is a slight departure from previously published techniques for *Drosophila* tissues^9,16,22^ in that explanted midguts are transferred to evenly spaced thin agarose pads and then covered with an additional layer of agarose (Figure 1O). This sandwiched structure allows the guts to be held in place for imaging, while also protecting them from damage that can result from their contact with the culture chamber walls if left freely floating. The agarose pads can be reproducibly cast and are thin enough (~100µm) not to interfere with imaging. Moreover, each agarose pad can comfortably house up to 3 midguts, allowing the multiple imaging of several explanted intestines. The agarose pads have the additional function of elevating the midguts from the bottom of the imaging chamber so that the surface of the agarose structure is directly exposed to air, creating an air-liquid interface. This is a key design element of the setup, as proper oxygenation was found to be essential for the long-term survival of explanted midguts (data not shown), similarly to what had been previously observed for the culture of wing imaginal discs^11^.

To increase the stability of the setup, agarose bridges connect each agarose sandwich (Figure 1O, middle panel), allowing the sample to survive 3 days of continuous imaging (Figure 2C and Video 2). Moreover, using a microscope equipped with an incubation chamber to control evaporation removes the need to replenish the imaging vessel with fresh culture medium. As such, the culture system, despite its simple design, is highly efficient and well suited for long-term imaging experiments.

**Figure 2:**
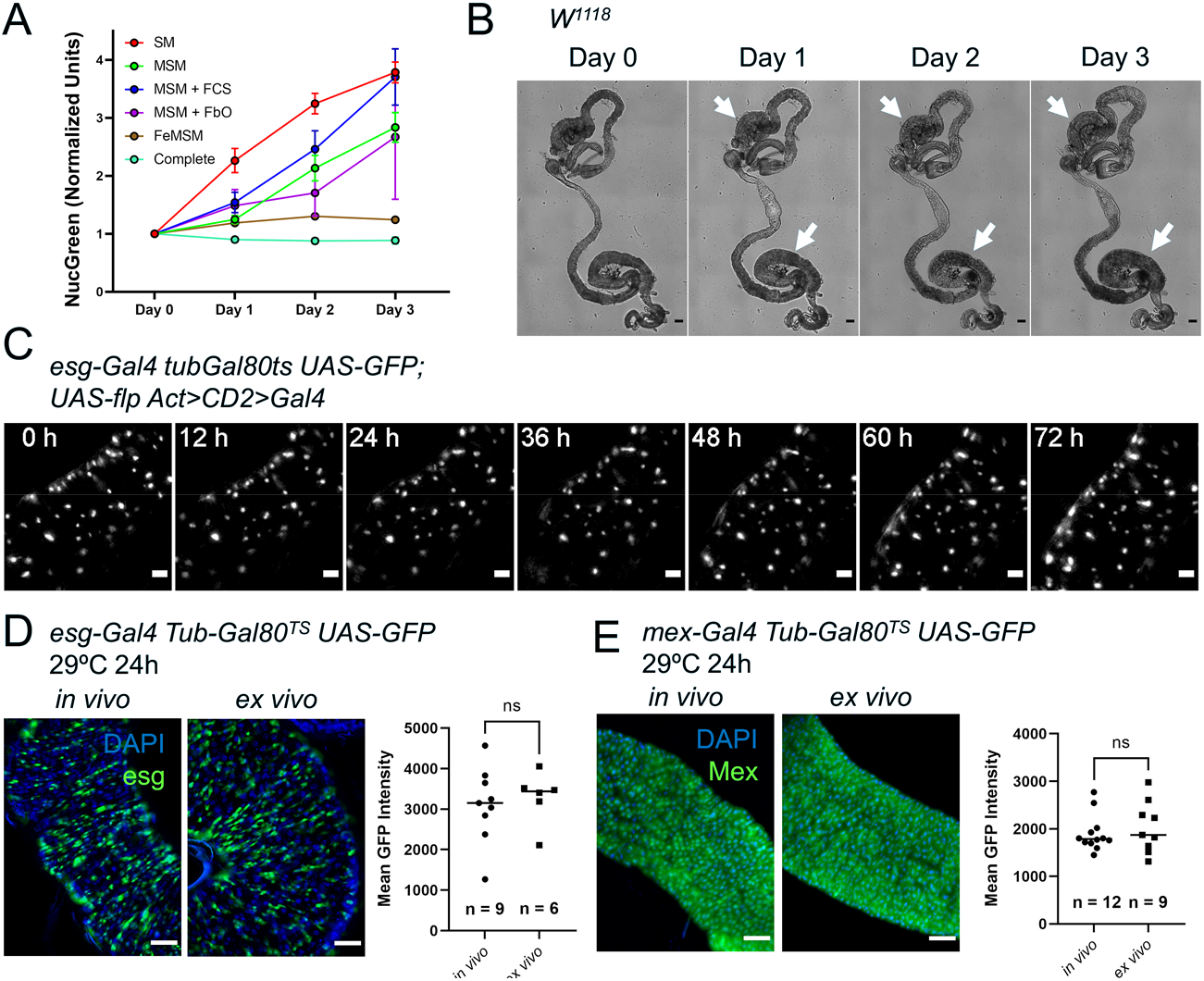
A custom culture medium sustains the midgut *ex vivo*. (A) Incorporation of the cell-impermeable dye NucGreen shows the levels of midgut cell death *ex vivo* in midguts cultured in: standard Schneider’s medium (SM), modified Schneider’s medium (MSM), MSM with 10% added fetal calf serum (FCS), MSM including co-culture with fat bodies and ovaries (FbO), culture in fly extract prepared in MSM (FeMSM), and the combination of all these conditions (Complete). Error bars represent the standard error of the mean. (B) Explanted midguts maintain their shape and tissue integrity for 3 days in culture. However, commensal bacteria keep growing in the lumen, especially in the posterior section where this can be seen as a darkening of the lumen (white arrows). Scale bar is 100µm. (C) Maximum intensity projection of intestine expressing a lineage-tracing system under *esg*^*TS*^ driver and imaged for 72h at 29°C. The fly of origin was incubated at 29°C for 24h prior to dissection. The intestine was undamaged during the course of imaging and did not show proliferation events. Scale bar is 5µm. See also Video 2. (D-E) Temperature-sensitive gene induction *ex vivo* is possible in both progenitor cells (D) and enterocytes (E). GFP levels expressed after 24h of gene induction are comparable between *in vivo* and *ex vivo* intestines. Explanted intestines were shifted to 29°C immediately after sample preparation. *In vivo* controls were shifted simultaneously. Images are representative maximum intensity projections of posterior midguts. Scale bar is 50µm. Each dot in the graphs represents the average GFP expression level in the posterior section of an intestine (T test). (ns, not significant)

### A culture medium tailored to adult *Drosophila* tissues enhances the survival of explanted midguts

One of the factors currently limiting the extended survival of explanted adult *Drosophila* tissues is the lack of culture media specifically designed for this task. To obviate this issue, we analyzed several parameters that distinguish the hemolymph of larvae, on which most current *Drosophila* cell culture media are based, to that of adult flies. We therefore compared the performance of different media formulations using the incorporation *ex vivo* of the cell-impermeable dye NucGreen (Thermofisher) as a measure of cell death (Figure 2A). The dye was added at the start of the culture, and its accumulation in the tissue measured over the course of three days. We found that raising the concentration of trehalose, which is found at high levels in *Drosophila* circulation^23–27^, and mimicking the Na^+^/K^+^ ratio of adult hemolymph^28–32^ is sufficient to significantly reduce cell death after 24h of culture. As Schneider’s medium^33–36^ is a common solution for several published protocols for the live-imaging of adult *Drosophila* tissues^14,16,17,19,21^, we modified it to raise the trehalose concentration to 50mM and Na^+^/K^+^ ratio to levels similar to those found in adult *Drosophila* hemolymph (Table 1)^24–26,28–32^. Our tests showed that supplementing the culture medium with 10% fetal bovine serum (FBS) did not have a significant beneficial effect on the midgut epithelium (Figure 2A), but we did notice that FBS addition resulted in the visceral muscle remaining intact and capable of peristalsis for longer (data not shown). Not surprisingly^37,38^, co-culturing explanted midguts with ovaries and abdomens lined with fat body (adipocytes) could also decrease cell death *ex vivo*. This effect was not due to the sequestration of NucGreen by ovaries and adipocytes as the dye was supplemented at a saturating concentration and its incorporation in these organs/tissues was mostly limited to areas mechanically damaged during dissection. As ovaries and fat bodies may enhance the survival of explanted midguts by secreting growth factors and/or nutrients, we reasoned that fly extract might have a similar effect. Indeed, midguts cultured in 100% fly extract prepared using modified Schneider’s medium had greatly reduced rates of cell death over time (Figure 2A). Lastly, as we designed the medium for the purpose of extended live-imaging, we also added N-acetyl cysteine and sodium citrate because their antioxidant effects reduced phototoxicity during extended imaging (data not shown). Combining all the findings mentioned above resulted in culture conditions (see Methods) that minimized cell death and allowed the live-imaging of explanted midguts for up to three days (Figure 2C and Video 2), a significant improvement over previously published culture protocols for the midgut in which imaging was limited to 16h^21,22^.

**Table 1:** Modified Schneider’s medium formulation. Components whose concentration was modified are highlighted in yellow.

| Component | Schneider's medium | Modified Schneider's medium |
| --- | --- | --- |
| Amino Acids | Concentration (mM) | Concentration (mM) |
| Glycine | 3.3 | 3.3 |
| L-Arginine | 2.3 | 2.3 |
| L-Aspartic acid | 3.0 | 3.0 |
| L-Cysteine | 0.5 | 0.5 |
| L-Cystine | 0.4 | 0.4 |
| L-Glutamic Acid | 5.4 | 5.4 |
| L-Glutamine | 12.3 | 12.3 |
| L-Histidine | 2.6 | 2.6 |
| L-Isoleucine | 1.1 | 1.1 |
| L-Leucine | 1.1 | 1.1 |
| L-Lysine hydrochloride | 9.0 | 9.0 |
| L-Methionine | 5.4 | 5.4 |
| L-Phenylalanine | 0.9 | 0.9 |
| L-Proline | 14.8 | 14.8 |
| L-Serine | 2.4 | 2.4 |
| L-Threonine | 2.9 | 2.9 |
| L-Tryptophan | 0.5 | 0.5 |
| L-Tyrosine | 2.8 | 2.8 |
| L-Valine | 2.6 | 2.6 |
| beta-Alanine | 5.6 | 5.6 |
| Inorganic Salts |  |  |
| Calcium Chloride (CaCl<sub>2</sub>) (anhyd.) | 5.4 | 5.4 |
| MgCl<sub>2</sub> Hexahydrate | 0.0 | 0.0 |
| Magnesium Sulfate (MgSO<sub>4</sub>) (anhyd.) | 15.1 | 15.1 |
| Potassium Chloride (KCl) | 21.3 | 21.3 |
| Potassium Phosphate monobasic (KH<sub>2</sub>PO<sub>4</sub>) | 3.3 | 3.3 |
| Sodium Bicarbonate (NaHCO<sub>3</sub>) | 4.8 | 4.8 |
| Sodium Chloride (NaCl) | 36.2 | 91.2 |
| Sodium Phosphate dibasic (Na<sub>2</sub>HPO<sub>4</sub>) anhydro | 4.9 | 4.9 |
| Other Components |  |  |
| N-Acetyl Cysteine | 0.0 | 2.0 |
| Trisodium Citrate Dihydrate | 0.0 | 1.0 |
| Alpha-Ketoglutaric acid | 1.4 | 1.4 |
| D-Glucose (Dextrose) | 11.1 | 11.1 |
| Fumaric acid | 0.9 | 0.9 |
| Malic acid | 0.7 | 0.7 |
| Succinic acid | 0.8 | 0.8 |
| Trehalose | 5.8 | 55.8 |
| Yeastolate (g/l) | 2000.0 | 2000.0 |

In examining cultured midguts, we observed the accumulation of luminal contents (*i*.*e*. previously ingested food) in the posterior midgut. This appeared to be caused by peristaltic movements of the visceral muscle (Figure 2B, white arrows). Peristaltic movements persisted despite the use of the calcium blocker isradipine, although this drug suppressed them significantly. These areas were found to darken and expand over the course of culture, an effect we attribute to the growth of enteric bacteria, which eventually caused cell death and tissue damage. We found that feeding flies fresh food in the days prior to an experiment and a short (~3-6h) starvation prior to dissection reduced the negative effect of food accumulation and growth of enteric bacteria. Supplementing the culture medium with antibiotics also enhanced explanted midgut survival. Lastly, it is reasonable to assume that axenically reared flies should be immune to the issue of growing enteric bacteria.

### Transgenic gene expression in midgut explant culture

One of the most striking features of *Drosophila melanogaster* as a research model is the wide range of readily available genetic tools, allowing the cell-type-specific and temporally-controlled activation or suppression of expression of genes of interest. The possibility to use such genetic tools in a live-imaging setup is highly advantageous, but so far the temporal control of gene expression *ex vivo* has proven unfeasible^21^. To further assay the behavior of midgut cells in explanted organs, we tested the induction of UAS-GFP by cell type-specific, temperature-sensitive Gal4 drivers (Figure 2D-E)^39,40^. As expected, midguts explanted from flies grown at the restrictive temperature (18°C) did not show any GFP expression (Video 11 and Figure 2 – figure supplement 1A, left panels). However, when incubated at the permissive temperature (29°C) at 0h after explant (Video 11 and Figure 2 – figure supplement 1A, right panels), they started expressing the UAS-GFP transgene. Moreover, GFP expression could also be induced in intestines cultured at 18°C for 24h and then shifted to the permissive temperature (29°C), indicating that the epithelium remains viable and genetically functional long-term (Figure 2 – figure supplement 1A). Indeed, the GFP intensity in midguts cultured at 29°C for 24h from the time of dissection was similar between *in vivo* and *ex vivo* conditions for both progenitor cells (Figure 2D) and enterocytes (Figure 2E) using the *escargot*-(*esg*) and *mex*-Gal4 Gal80^TS^ driver lines, respectively. This indicates that transcription and protein synthesis are maintained at normal levels in our culture system and shows that this system can be used to assay the effects of transgene induction in real time. Interestingly, progenitor cells were found to be asynchronous in their expression of the reporter GFP (Fig 4A and Video 11), with some starting to reach detectable levels only 20 hours after the first GFP^+^ cells become visible. This could be explained by variations in the activity of the *esg* promoter or by different global rates of transcription and translation, which in turn could be indicative of different cell states.

### Progenitor cells require stimulation to proliferate and differentiate

When imaging explanted adult *Drosophila* midguts we observed *esg*-expressing progenitor cells (ISCs and EBS) to be quiescent (Figure 3A and Videos 2 and 3). Progenitor cells in explanted healthy guts did not show changes in their GFP levels, indicative of continuous *esg* expression, nor in their nuclear size (Fig 2D), which indicates that DNA content remains constant. Via cell tracking we also observed that GFP-labeled progenitor cells did not divide in healthy explanted intestines (Figure 4D). It is important to note that in said healthy intestines we did not observe cell death or enterocyte (EC) extrusion events until after 48-72h of culture. As healthy enterocytes are known to suppress ISC proliferation^41^, this may explain the lack of cell division in our explants. The suppression of peristaltic movements by isradipine may also prevent EC loss by reducing mechanical tissue stress^42^.

**Figure 3:**
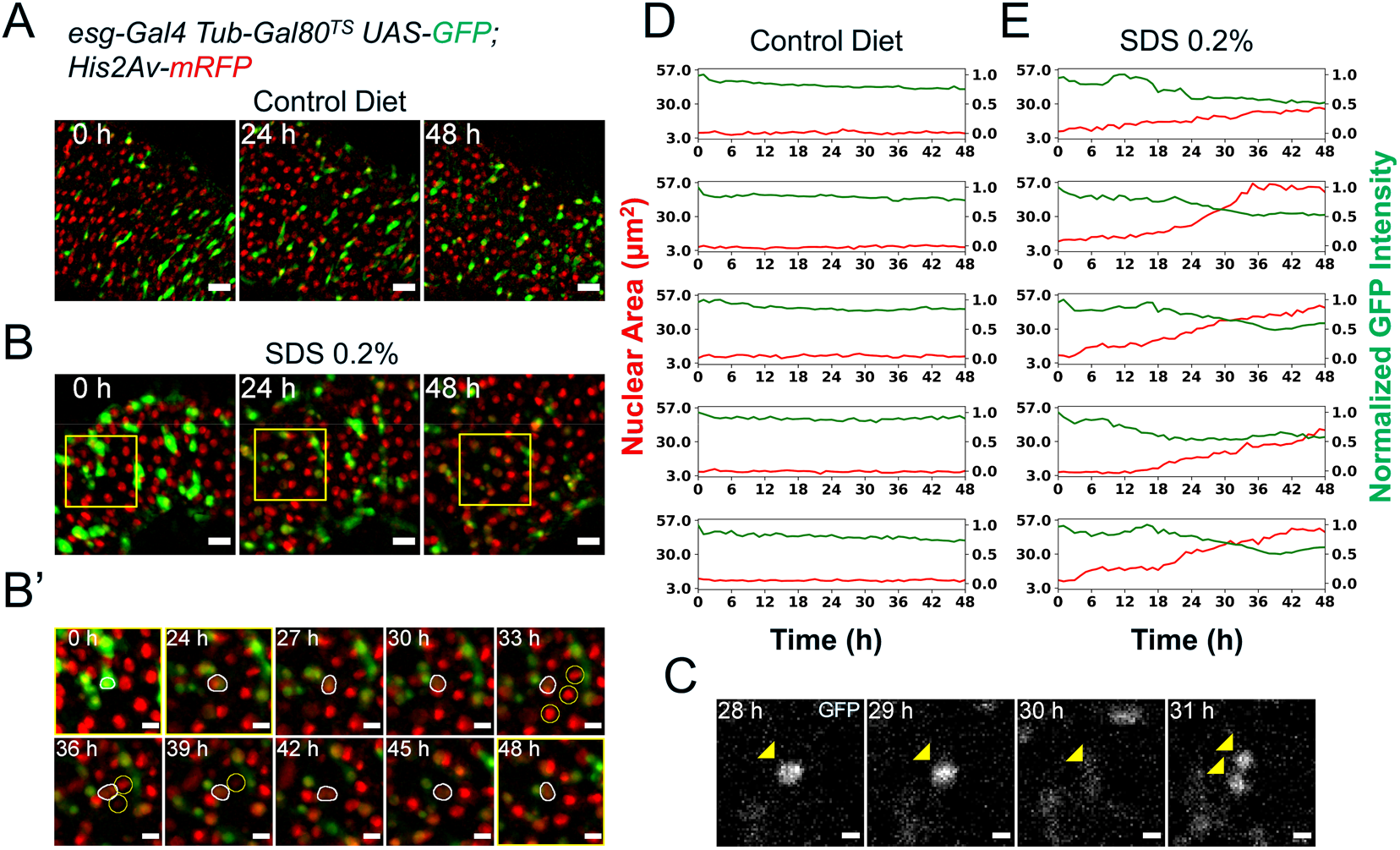
Midguts respond to damage *ex vivo*. (A) Healthy gut fed a control diet showing no sign of tissue damage. Images are maximum intensity projections. See also Video 3. Scale bar is 20µm. (B) Gut from fly fed SDS 0.2% overnight showing progressive tissue repair *ex vivo* mediated by progenitor cells proliferation and differentiation. As cells differentiate, GFP expression is gradually lost. Area delimited by the yellow rectangle is enlarged in (B’). Images are maximum intensity projections. See also Video 4. Scale bar is 20µm. (B’) Enlargement of panel (B) showing an *esg*^+^ cell (white outline) differentiating and replacing dying enterocytes (yellow circles). As the cell differentiates, GFP expression is lost and its nucleus grows larger. Images are maximum intensity projections. See also Video 5. Scale bar is 10µm. (C) Example of stem cell dividing upon tissue damage initiated by SDS feeding. Cell is marked by the expression of nuclear GFP (yellow arrowhead). Note that in the 30h time-point the cell is in mitosis and so the nuclear GFP signal is mostly lost due to nuclear envelope breakdown. After mitosis, the nuclear envelope is reformed and GFP re-accumulates in the nucleus of the daughter cells. Images are single z-slices. Scale bar is 5µm. (D) Plots of nuclear area (red) and mean nuclear GFP intensity (green) from single progenitor cells from healthy guts. Note that both nuclear size and GFP signal, despite a small gradual dip caused by photobleaching, are stable for the duration of the imaging session. (E) Plots of nuclear area (red) and mean nuclear GFP intensity (green) in single progenitor cells from SDS-damaged guts. Note that nuclear size increases, while GFP signals dims over time, suggesting EB to EC differentiation.

**Figure 4:**
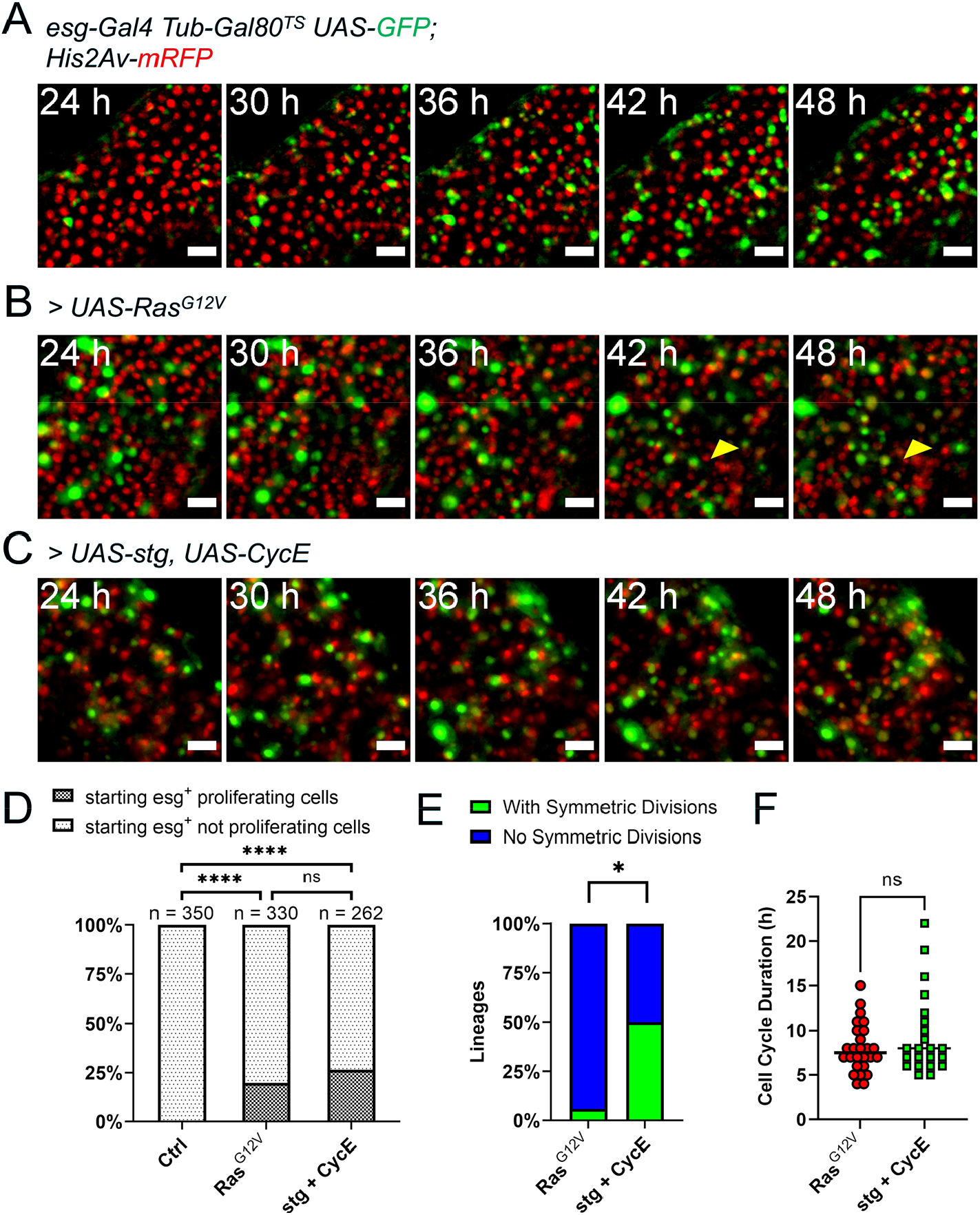
Genetic induction of intestinal stem cell proliferation. (A) Live-imaging of a normal intestine expressing GFP in progenitor cells via *esg*^*TS*^ driver. Intestines were shifted to 29°C at the start of imaging. Note the gradual accumulation of GFP signal showing variability between cells in terms of both time and intensity. Images are maximum intensity projections. See also Video11. Scale bar is 20µm. (B) Live-imaging of an intestine expressing GFP and *Ras*^*G12V*^ in progenitor cells via *esg*^*TS*^ driver. Intestines were shifted to 29°C at the start of imaging. GFP^+^ cells can be seen rapidly dividing and displacing mature enterocytes (yellow arrowhead). Images are maximum intensity projections. See also Video12. Scale bar is 20µm. (C) Live-imaging of an intestine expressing GFP and *Ras*^*G12V*^ in progenitor cells via *esg*^*TS*^ driver. Intestines were shifted to 29°C at the start of imaging. GFP^+^ cells can be seen rapidly dividing. Images are maximum intensity projections. See also Video 13. Scale bar is 20µm. (D) Quantification of observed proliferating and non-proliferating esg^+^ progenitor cells in control and intestines expressing Ras^G12V^ or stg and CycE. Only cells observed from the moment they expressed visible levels of GFP to the end of the imaging session are included. The progeny of observed mitoses were not counted for this analysis. (E) Frequency of stem cell lineages with observed symmetric mitoses, compared between intestines expressing either *Ras*^*G12V*^ or *stg* and *CycE* (Fisher’s exact test). (F) Quantification of cell cycle durations of progenitor cells expressing either *Ras*^*G12V*^ or *stg* and *CycE*, and GFP. No significant difference was found between the two genotypes (Mann-Whitney test). (ns, not significant; *, p < 0.05)

A key feature of *Drosophila* intestinal progenitor cells is their ability to rapidly respond to tissue damage^1–3,5,43^. To confirm that this capability is maintained *ex vivo*, we fed female flies Sodium dodecyl sulfate (SDS) mixed with solid fly food at a final concentration of 0.2% (v/w) for 18h. Following this treatment, the flies were fed 0.05% sucrose in aqueous solution on a cotton pad for an additional 4-6h prior to dissection, in order to flush the SDS from the gut. This protocol caused only mild initial tissue damage, and was advantageous as it did not result in the retention of high levels of SDS in explanted guts. However, by mixing the SDS-laced food with a blue food-safe dye, we found that some low amounts of SDS-laced food do persist in the lumen of intestines at the time of dissection. The transient exposure to SDS and the low amounts retained in the lumen resulted in gradual EC death and/or extrusion, allowing the live-imaging of damage response. When imaged, intestines from SDS-fed flies showed progenitor cell proliferation events (Figure 3C). However, ISC mitoses could generally be observed only in cases where tissue damage was extensive, as evidenced by the appearance of pyknotic or fragmented nuclei and the extrusion of multiple (> 3) contiguous ECs. As this tended to happen towards late time-points after dissection, only a few (1-2) mitotic events per imaged field could be observed with this damage protocol. In addition to ISC divisions, many progenitor cells could be observed growing in nuclear size while simultaneously losing GFP expression (Figure 3B,E and Videos 4-5), suggestive of differentiation events towards the EC identity. Indeed, differentiating progenitor cells could be seen replacing dying enterocytes (Video 5).

A limitation of the SDS-feeding protocol described above is that midguts were damaged prior to imaging. Therefore, it cannot be excluded that the observed differentiation events could be due to EBs already primed to differentiate during the time of feeding. To confirm that, at the time of damage, pre-existing EBs were indeed capable of responding to tissue stress, we used a thin tungsten needle to create a small lesion in explanted intestines. The lesions perforated both visceral muscle and epithelial layers, but left the peritrophic matrix intact (Figure 3 – figure supplement 1A). Intestines were then enveloped in agarose and imaged within 20’ of the time of damage. Similarly to the SDS damage protocol, we were able to record progenitor cells both dividing and differentiating (Figure 3 – figure supplement 1B,C and Video 6). Moreover, wounding stimulated the robust expression of the cytokine Unpaired 3 (Upd3), whose function in midgut damage response is well documented (Figure 3 – figure supplement 1D,E and Video 8)^3,44,45^.

All in all, these observations suggest that both ISCs and EBs are quiescent in undamaged intestines. While previous studies did show varying rates of ISC proliferation in healthy midguts *in vivo*^1–3,46,47^, this may be attributed to cell death^41^ as well as the passage of food through the intestinal lumen^42^, both of which do not occur in our midgut cultures until later time-points (*i*.*e*. after 48-72h). Moreover, many previous studies used lineage-tracing tools requiring 37°C heat shock for their activation, a treatment known to increase ISCs proliferation rates^4,47–50^. Indeed, when in the presence of tissue damage progenitor cells could be robustly activated in explanted intestines, indicating that both ISCs and EBs were still capable of responding to their environment *ex vivo*.

### Loss of *Notch* drives tumorigenesis *ex vivo*

A key pathway necessary for progenitor cell differentiation is the *Delta*/Notch signaling that occurs between ISCs and adjacent EBs^1,2,51^. The depletion of *Notch* (N) by RNA interference (*N*^*RNAi*^) in esg^+^ cells has been shown to promote the formation of undifferentiated tumor-like masses *in vivo*^1,2,44,51^. Notably, tumor initiation by N-depleted ISCs requires proliferation induced by tissue stress^44,52^. Consistent with this, when we explanted midguts from flies expressing N^RNAi^ in progenitor cells for 24h prior to dissection and cultured them for 48h, we did not observe tumorigenesis in the absence of tissue damage (Figure 3 – figure supplement 2A and Video 9). However, when guts were accidentally damaged during dissection (Figure 3 – figure supplement 2B, yellow ellipses), esg^+^ cells started to proliferate (Figure 3 – figure supplement 2B, yellow squares, and Video 10). Notably, progenitor cells remained small and did not lose GFP expression (compare Figure 3B and Figure 3 – figure supplement 2B as well as Videos 3 and 10), suggesting a block of differentiation *ex vivo* in response to *Notch* knock-down.

### Intestinal progenitor proliferation can be genetically stimulated *ex vivo*

EGFR-Ras-Erk signaling is activated upon gut tissue damage, and this pathway is required for stem cell activation in the adult *Drosophila* intestine^53–55^. Previous experiments showed that expression of a constitutively active form of *Ras* (*Ras*^*G12V*^) strongly promotes stem cell proliferation in adult midguts^55,56^. As our culture protocol allows the expression of transgenes *ex vivo*, we induced *Ras*^*G12V*^ in explanted midguts using the *esg-Gal4 Gal80*^*TS*^ progenitor-specific driver gene combination (*esg*^*TS*^). When explanted midguts were shifted to the permissive temperature (29°C) at the start of imaging, progenitor cells, marked by GFP co-expressed with *Ras*^*G12V*^, started to rapidly proliferate (Figure 4B). About 20% of progenitor cells tracked from the moment they expressed visible amounts of GFP till the end of the imaging session, were seen proliferating (Figure 4D). The progeny of observed mitoses were not counted for this analysis. However, since many cells were not trackable for the duration of the imaging session due to major tissue rearrangements, we may be underscoring the percentage of proliferative GFP^+^ cells. As GFP^+^ cells accumulated in the tissue, enterocytes were displaced and extruded from the epithelium (Figure 4B, yellow arrowhead, and Video 12). Interestingly, some GFP^+^ progenitor cells did not proliferate, but their nuclei rapidly grew in size, which aligns with the previously reported role of EGFR signaling in promoting EB growth^55,57^. Moreover, these rapidly growing progenitors also lost GFP expression during the course of imaging, which suggests their differentiation towards the EC lineage.

The extended live-imaging that our protocol allows also permitted us to follow cells through multiple rounds of mitosis. Using manual 3-dimensional (3D) cell tracking, we reconstructed the lineages of 17 dividing cells that expressed *Ras*^*G12V*^ (see Figure 5 and Video 14 for an example). As ISCs are the only cell type in the *Drosophila* intestine capable of dividing multiple times^1,2,58^, the founding cells in these lineages were *bona fide* ISCs. We directly measured the duration of the ISC cell cycle in these lineages at 8 ± 2.76h (Figure 4F). Interestingly, of 17 *Ras*^*G12V*^-expressing ISC lineages characterized, only one had clearly symmetric mitoses (Figure 4E), defined as division events that gave rise to daughter cells which subsequently both divided again (Figure 6A-B). This is in accordance with previously published data showing that most mitotic events in the adult intestine are asymmetric^1,2,47^, producing one new ISC and one cell that differentiates towards the EC or enteroendrocrine (EE) cell fate, as well as the observation that Ras^G12V^ does not prevent ISC differentiation as previously reported^55^.

**Figure 5:**
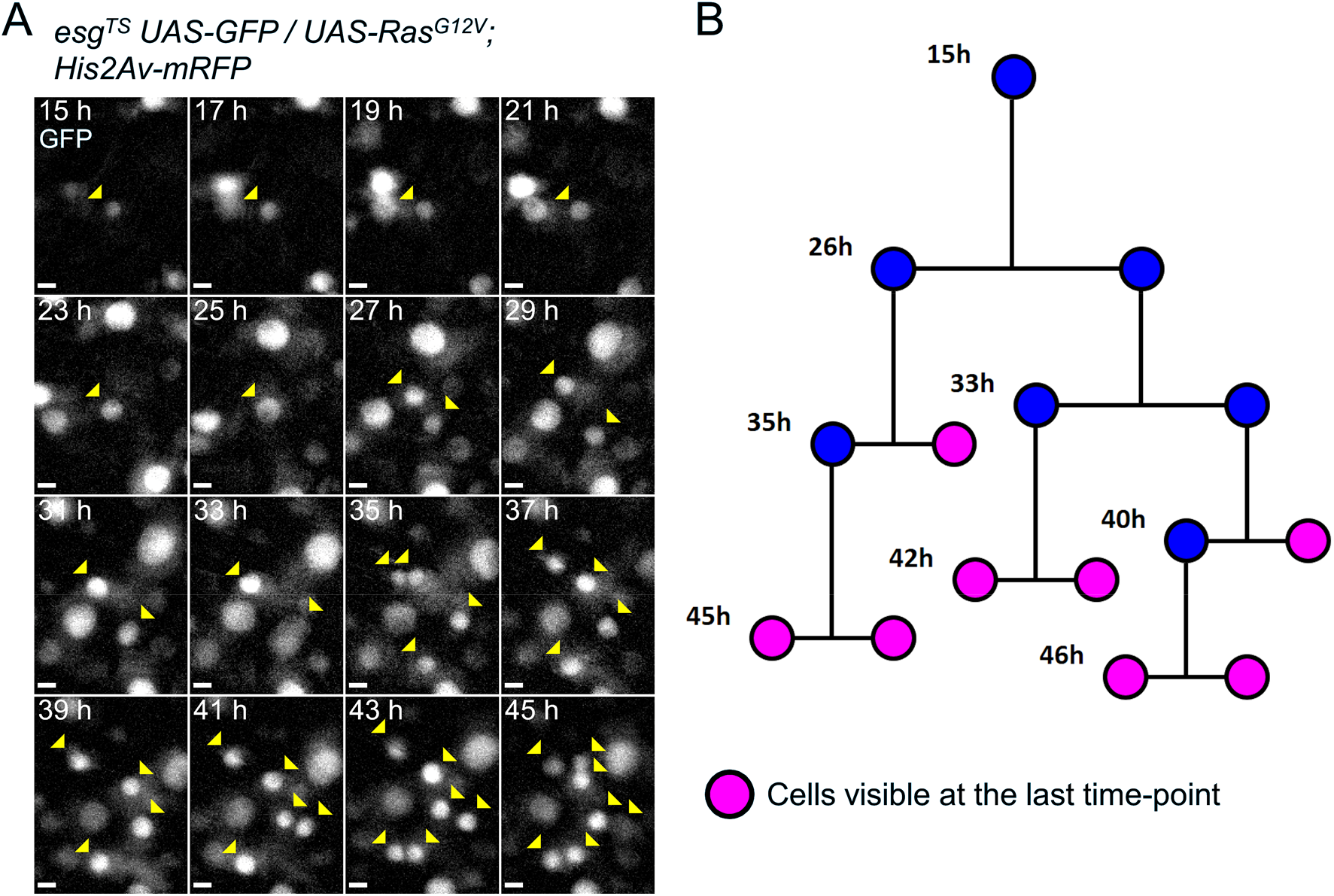
Example of a reconstructed ISC lineage. (A) Time-lapse of a single Ras^G12V^-expressing ISC undergoing multiple rounds of mitoses. The GFP channel is shown. Cells belonging to the lineage are marked by yellow arrowheads. (B) Lineage diagram. Cells visible at the last time-point shown are marked in magenta. Images are maximum intensity projections of z-slices encompassing the cells in the lineage. See also Video 14. Scale bar is 5µm.

**Figure 6:**
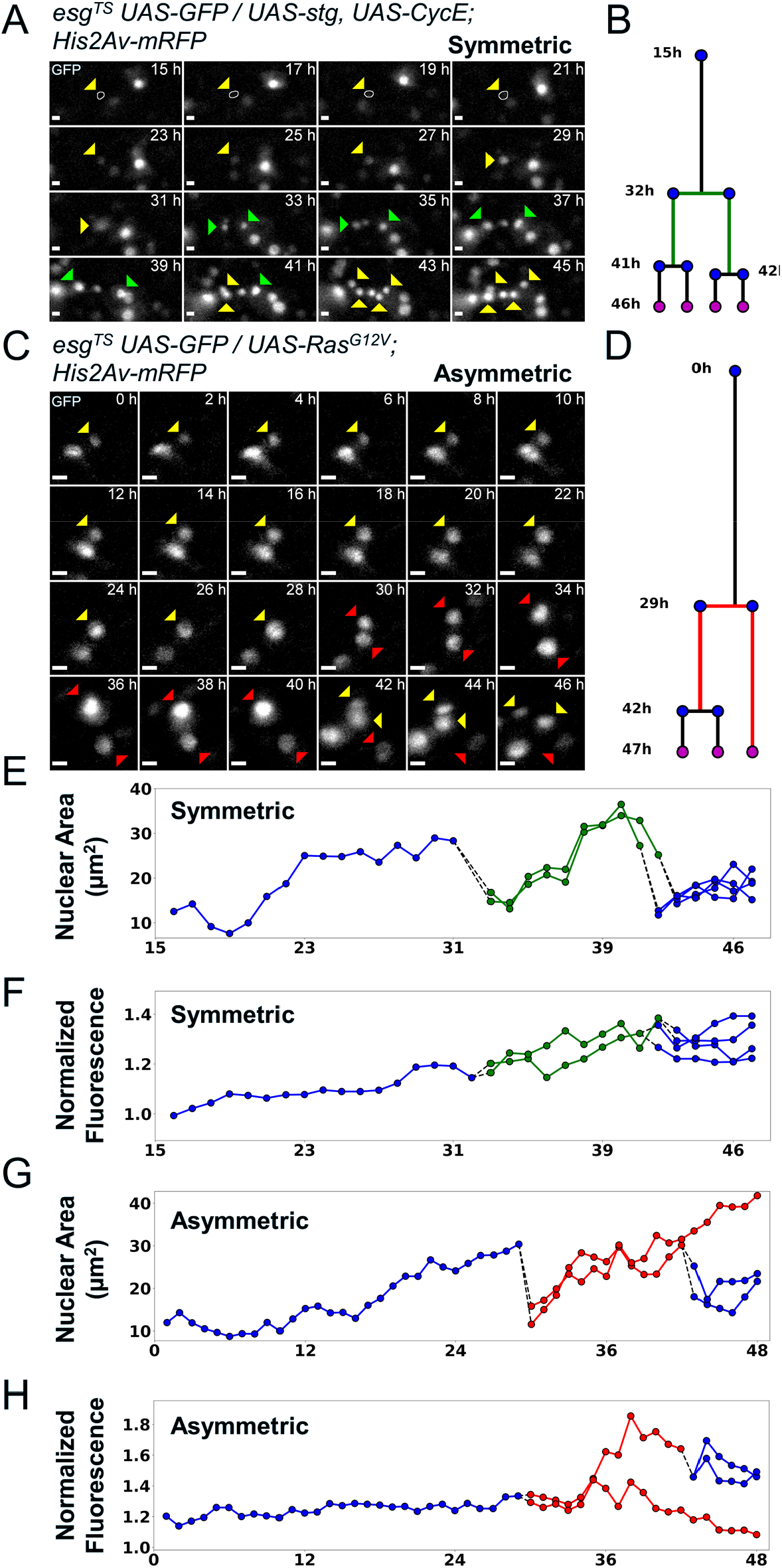
Examples of asymmetric and symmetric divisions. (A) GFP channel showing a stem cell undergoing symmetric division upon expression of *stg* and *CycE* via *esg*^*TS*^ driver. The resulting daughter cells (green arrowheads) can be seen further dividing at the 41h and 42h time-points. A white circle denotes the outline of the starting cell at the initial time-points, when signal was weakest.

Since *Ras*^*G12V*^ stimulation could have a role in cell differentiation, we tested whether stimulating ISC proliferation more directly would result in more symmetric lineages. For this we used the *esg*^*TS*^ driver to express *string* (*stg*), a *Cdc25C* homolog, and *Cyclin E* (*CycE*) (Figure 4C and Video 13). *String* directly activates *Cdk1* to promote mitosis, whereas *CycE* directly activates *Cdk2* to promote DNA replication and S-phase progression. The combined expression of these two gene products is sufficient to strongly induce ISC proliferation^59^. The fraction of progenitor cells that divided in response to Stg and CycE co-expression (~25%; Figure 4D) was not significantly different from that observed after forced expression of Ras^G12V^, suggesting that all receptive ISCs are activate in both cases. However, symmetric mitoses appeared in 6 of 12 ISC lineages that overexpressed *stg* and *CycE* (Figure 4E), a significantly higher frequency to that observed after *Ras*^*G12V*^ overexpression (p = 0.0106, Fisher’s exact test). While this difference might be explained by faster cell cycling due to *stg* and *CycE* promoting cell cycle progression more directly than Ras^G12V^, in fact we detected no significant difference in cell cycle duration in ISC cell cycles driven by *stg* and *CycE* (9.4 ± 4.6h) and cell cycles driven by *Ras*^*G12V*^ (8 ± 2.76h; Figure 4F).

### The progeny of symmetric mitoses actively move apart

Combining all the lineages described above, we were able to identify 8 symmetric divisions (see Figure 6A-B and Video 15 for an example). For 7 of these, the two daughter cells divided within 2h of each other, while for one pair the interval was 5h. This suggests that, in most cases, the sister cells originating from symmetric divisions will have cell cycles of similar length. Based on this, we defined divisions as functionally asymmetric if only one daughter cell was seen dividing further (see Figure 6C-D and Video 16). Further, to define a lineage as asymmetric, the non-dividing sister cell needed to remain quiescent for the remainder of the imaging session, which had to last at least 6h after the sister’s mitotic event (*i*.*e*. the second division in the lineage) (Figure 6D). Following this definition, we classified 17 sister pairs as asymmetric (see Figure 5D-F for an example), 9 from *Ras*^*G12V*^- and 8 from *stg* and *CycE*-expressing guts. Interestingly, for 8 of these 17 pairs, the non-dividing cells displayed increases in nuclear size over time, and also lost GFP expression, which is indicative of differentiation towards the EC cell fate (Figure 6C,G-H). This behavior was not observed for symmetric pairs, where both cells increased their nuclear size before dividing further, but maintained GFP expression (Figure 6A,E-F).

It was previously reported that spindle orientation is indicative of whether an ISC mitosis is asymmetric or symmetric, with the latter being essentially planar relative to the visceral muscle layer^22,51,60^. As we restricted the interval between subsequent imaging frames to 1h to reduce phototoxicity, we were not able to measure spindle orientations. However, it has been proposed that the orientation of the mitotic spindle results in daughter cells residing at different levels within the pseudostratified intestinal epithelium, with new-born EBs being more apical^51^. We therefore selected mitotic events that occurred in regions of epithelium that were flat with reference to the imaging plane (based on cell nuclei positions). The 3D profile of sister cells after mitosis was then reconstructed and we measured the angles between sister cells. Measured angles were found to be <15° in 29/33 cases for *Ras*^*G12V*^- and in 30/32 cases for *stg*+*CycE*-induced mitoses. Moreover, no significant differences in sister cell angles were observed between asymmetric and symmetric lineages (Figure 7A-C). This suggests that daughter cells, regardless of their eventual fates, may rearrange themselves after the mitotic event to sit basally on the basement membrane. Indeed, a previous study by Martin *et al*^21^ showed that even mitotic spindles can re-orient several times during the course of mitosis.

**Figure 7:**
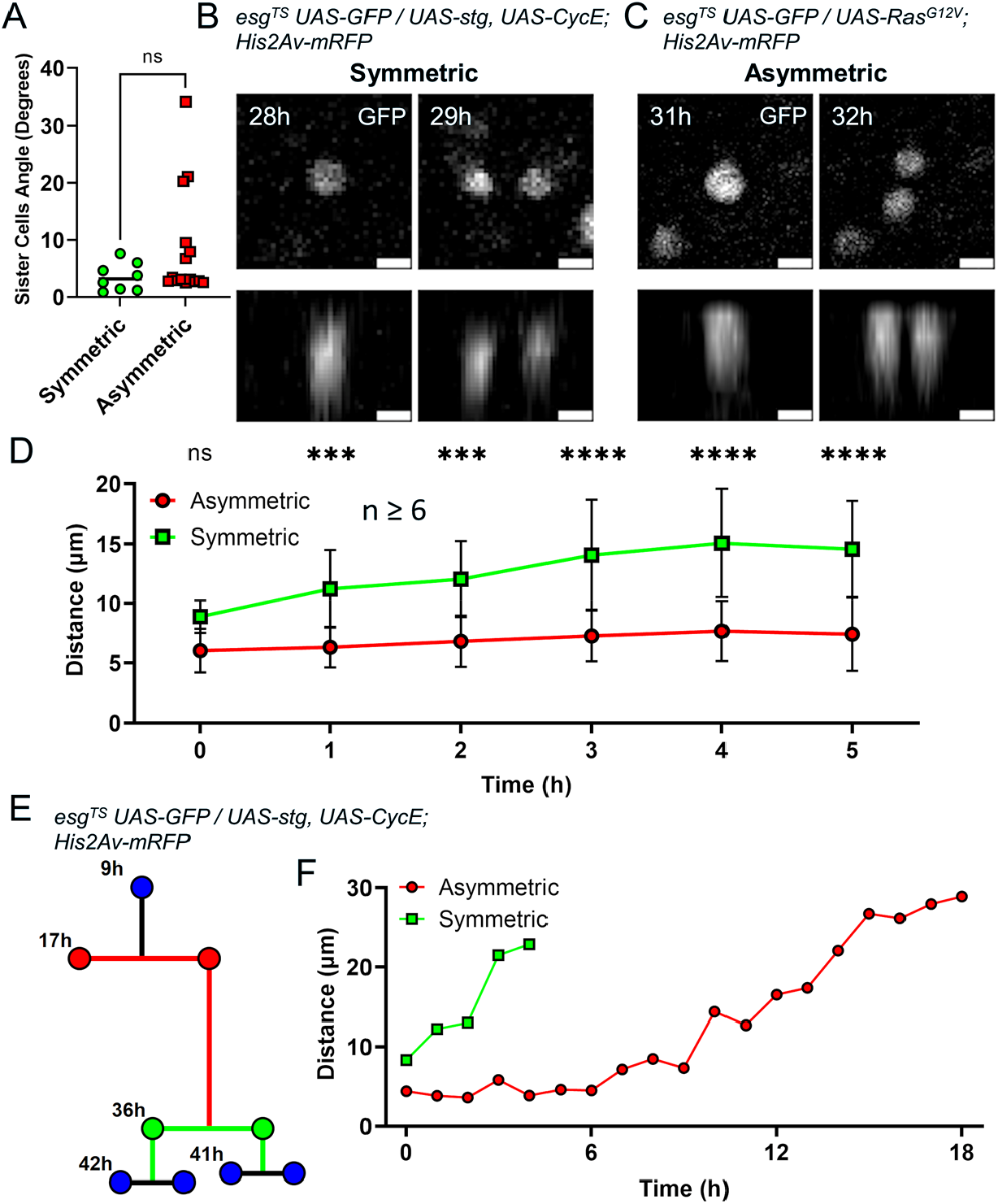
Analysis of asymmetric and symmetric divisions in the *Drosophila* midgut. (A) Angle between daughter cells after mitosis as referenced to the imaging plane. No difference was found between the progeny of asymmetric and symmetric divisions (Mann-Whitney). (B) XY (top panel) and Z (bottom panels) GFP profiles of the symmetric division event from Figure 5A,B. Cell proliferation was stimulated by *esg*^*TS*^-driven expression of *stg* and *CycE*. Scale bar is 5µm. (C) XY (top panel) and Z (bottom panels) GFP profiles of the asymmetric division event from Figure 5C,D. Cell proliferation was stimulated by *esg*^*TS*^-driven expression of *Ras*^*G12V*^. Scale bar is 5µm. (D) Internuclear distance between daughter cells in the first 5h after mitosis. Time point 0h denotes the first at which the two daughter cells are visible. Error bars represent standard deviation (Two-way Anova and Šídák’s multiple comparisons test). (E) Example of lineage characterized by both an asymmetric division (red) and a symmetric one (green). Cell proliferation was stimulated by *esg*^*TS*^-driven expression of *stg* and *CycE*. (F) Internuclear distance between the daughter cells of the asymmetric and symmetric division from the lineage in E. Time point 0h denotes the first at which the two daughter cells are visible. (ns, not significant, ***, p < 0.001; ****, p < 0.0001)

A significant difference between the symmetric and asymmetric divisions was, however, found when we tracked newborn cells over time. Due to some cells having cell cycles lasting less than 8h, we only considered the first six 1h time-points following the appearance of a sister pair (defined as time-point 0).

Measuring the distance between sister cells at each time interval, we found that cells in asymmetric pairs remained close to one another until the following mitotic event (Figure 7D, red line). Cells in symmetric pairs, however, moved apart from one another (Figure 7D, green line). These behaviors could be observed even when considering the progeny of asymmetric and symmetric mitoses from the same lineage (Figure 7E-F). This observation suggests that commitment to differentiation, as occurs after asymmetric divisions, requires that sister cells remain in contact for ~3-5 hours (Figure 7D,F, red line). Conversely, rapid separation of sister cells following a division may be necessary to generate a symmetric division that duplicates ISCs.

Differences in the motility of daughter cells of asymmetric and symmetric divisions could explain these observed effects. We therefore measured cell motility as the distance in 3D space that a cell travelled between one time-point and the next. We found that the genotype used to induce proliferation did not have an effect on the cell motility of either the dividing or non-dividing progeny of asymmetric divisions (Figure 7 – figure supplement 1A). Likewise, no difference in cell motility was found when considering the effect of time on cell movement of the progeny of asymmetric (Figure 7 – figure supplement 1B,C) or symmetric (Figure 7 – figure supplement 1D) divisions. Therefore, we directly compared the motility of cells generated by asymmetric or asymmetric divisions irrespective of genotype or time-point analyzed (Figure 7 – figure supplement 1E) and found no significant difference. This indicates that the progeny of both symmetric and asymmetric divisions migrate within the epithelium at similar speed. Since the progeny of asymmetric divisions tend to remain close to one another over time, their movement is most likely random. On the other hand, since the distance between daughter cells derived from symmetric mitoses increases over time, their movement is directional, with the cells in the pair moving away from each other. It would therefore be expected that, when considering symmetric sister pairs, the relative movement of one cell to its sister should be greater than that observed for asymmetric pairs. We therefore considered the motion of sister pairs, decomposing their movement in X-and Y-axis components and summed the resulting vectors, thus calculating the movement of a cell relative to its sister along the X- and Y-axis. As expected, the magnitudes of the reconstructed relative movements between symmetric pairs was significantly greater than that for asymmetric pairs.

### *Ex vivo* culture of other adult *Drosophila* organs

To test the feasibility of our culture setup to sustain other adult *Drosophila* organs, we first focused on the Malpighian (renal) tubules. Being physically connected to the intestine, we reasoned the two organs may share similar requirements for their survival *ex vivo*. A shared characteristic between Malpighian tubules and the adult midgut is the presence of a population of progenitor cells marked by *esg* expression^61,62^. Using the *esg*^*TS*^ driver system, GFP expression could be induced in tubules cultured at 18°C for 24h and then shifted to the permissive temperature (29°C), indicating their long-term survival (Figure 8A-B). Moreover, we found that Malpighian tubules cultured for 3 days could still contract regularly (Video 17), albeit only if still attached to intestines.

**Figure 8.**
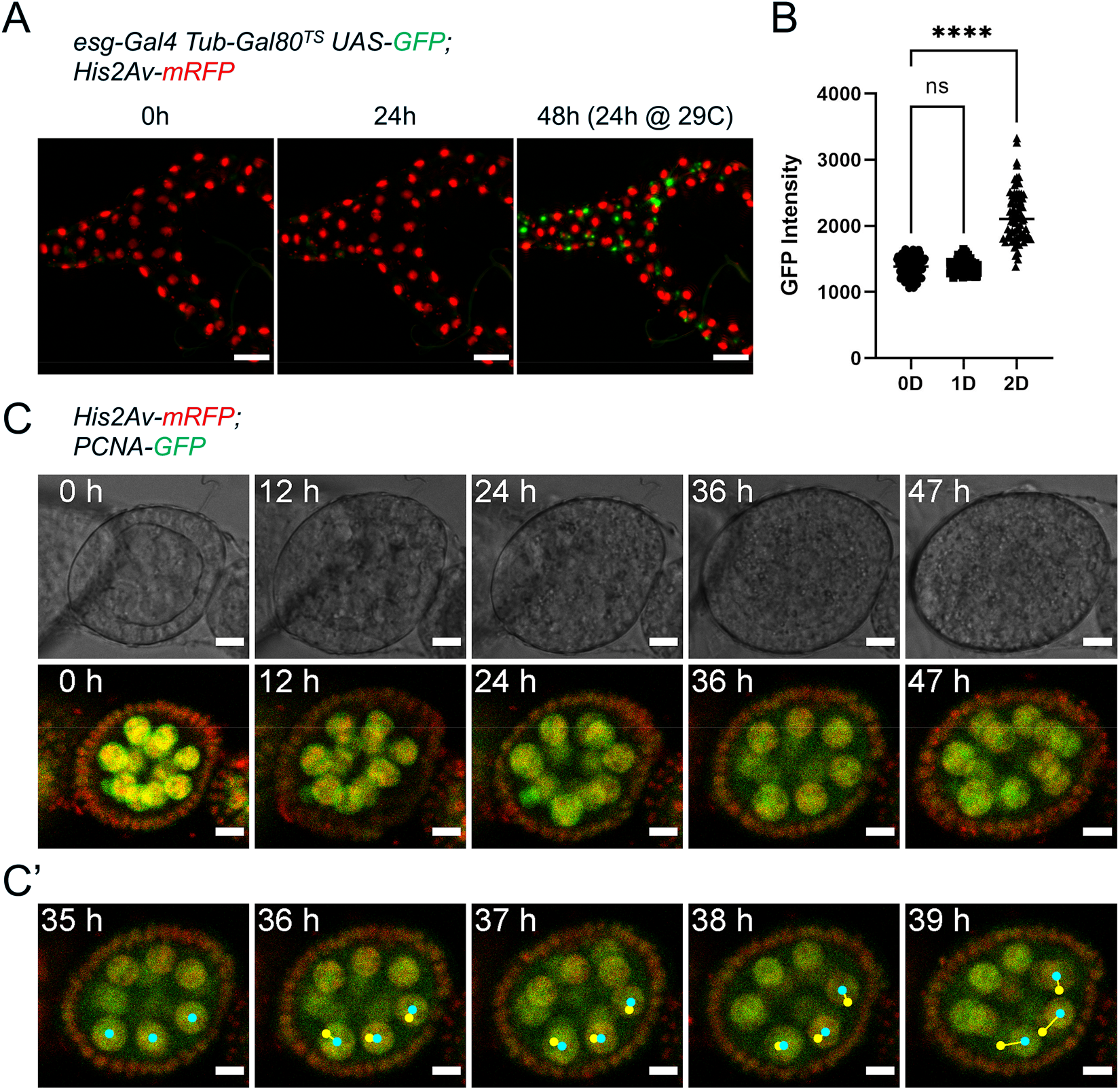
*Ex vivo* culture of Malpighian tubules and ovaries. (A) Malpighian tubules cultured at 18°C for 24h, then shifted to 29°C are still able to activate expression of GFP driven by the *esg*^*TS*^ system, showing that the epithelium remains healthy long-term. Images are maximum intensity projections. Scale bar is 50µm. (B) Quantification of GFP expression of progenitor cells from Malpighian tubules cultured *ex vivo*. Same cells were measured at 0, 1, and 2 days after explantation. (Dunnett’s multiple comparisons test) (ns, not significant, ****, p < 0.0001) (C) Stage 4 follicle growing in size and elongating over the course of 2 days *ex vivo*. (C’) Selected frames showing follicle rotation. Cyan dots mark the current position of a nucleus, while yellow ones marked the position in the previous frame. Images are maximum intensity projections of the 3 center-most z-slices. See also Video 20. Scale bar is 10µm.

A key component of our culture system that prolongs the viability of midguts is the co-culture with dissected abdomens and ovaries. On closer inspection, we found that adult hearts, laying along the abdominal cuticle, could be seen still beating regularly after up to 10 days in culture (Video 18).

Similarly, the muscle sheet that envelopes the ovaries was still contracting after 3 days in culture (Video 19). We then took a closer look at ovaries by dissecting individual ovarioles for live-imaging. A distinctive characteristic of stage 5-8 follicles is their rotation within their follicle cell sheath, along their long axis^63^. In our *ex vivo* cultures, we routinely observed stage 4 follicles growing in size and starting to rotate and elongate (Figure 8C and Video 20), indicative of progression to stage 5. Notably, this phenomenon continued for up to 48h, suggesting the long-term survival of follicles in our explants.

Therefore, we believe our culture protocol could be applied to other adult Drosophila tissue and could prove useful in investigating a wide range of biologically relevant phenomena.

## DISCUSSION

The adult *Drosophila* midgut has emerged as a powerful tool to understand the biology of epithelia and their resident stem cells. In recent years this system has been enriched by the development of advanced live-imaging approaches^18–22^ that allow the observation of adult midguts for up to 16h. However, many biologically interesting processes, for instances regeneration, occur over longer time-spans, and so methods for extended culture and live-imaging should prove advantageous.

Several factors could cause the limited survival of midguts *ex vivo*. Firstly, currently available *Drosophila* culture media are based on larval hemolymph composition. We have shown here that minimal modifications introduced to Schneider’s medium are sufficient to reduce cell death *ex vivo* (Figure 2A). Midguts may also receive nutrients and signaling molecules from other organs such as ovaries and fat body, which are in close proximity to the intestine. Indeed, using fly extract as a culture medium or co-culturing intestines with ovaries and fly abdomens resulted in a dramatic decrease in cell death (Figure 2A). Proper oxygenation is also a concern as it had been found to be essential for other *Drosophila* organ *ex vivo* cultures^11^. Indeed, trachea ramify throughout the fly’s internal organs, and in the intestine they even reach through the visceral muscle to contact epithelial cells directly^64^. Therefore, we designed our culture setup to keep guts elevated and close to the surface of the culture medium at a liquid-air interface (Figure 1). Moreover, the sample setup was designed to be efficient and simple to construct for ease of reproducibility, allowing up to 12 explanted midguts to be imaged in parallel in a single dish. Lastly, prolonged live-imaging sessions can be hampered by phototoxicity. This can be resolved by reducing the intensity of the excitation light and exposure times, albeit at the cost of a reduced signal/noise ratios and frame rates. Controlling each of these factors has allowed us to culture healthy explanted midguts for up to 3 days *ex vivo* (Figure 2C and Video 2), and other organs for even longer periods.

Using our system, we observed that, while midguts maintain their ability to respond to tissue stress *ex vivo*, progenitor cells in undamaged intestines are quiescent. Previous estimates of mitotic rates based on immunostaining for the mitotic marker phospho-Ser 10-Histone 3 generally showed a wide range of baseline values, with numbers even as low as 1-3 mitoses per midgut^3^. Nonetheless, even considering a low mitotic rate and using an estimate of mitosis duration^21^ and ISCs numbers^1,65–67^, we would have expected to see several mitoses even in undamaged intestines (*e*.*g*. >5 for fields with 50 or more *esg*^+^ cells). As ISC proliferation could be stimulated by tissue damage *ex vivo*, this suggests that, in homeostatic conditions, stem cells may only proliferate when the need to replace damaged or dying cells arises. Given the lack of cell death in undamaged midguts *ex vivo*, the previously reported proliferation-suppressive effect of enterocytes^41^ may be responsible for the lack of observed mitotic events. Interestingly, we also did not observed differentiation events in undamaged intestines, which suggest that enteroblasts are a stable cell type, rather than being transient progenitors that are present only during periods of rapid ISC division, and are rapidly lost via differentiation or apoptosis^46^. Indeed, a previous analysis of the EB gene expression profile did show the existence of EB-specific genes, consistent with EBs being a distinct cell type^68,69^. As a consequence, the enteroblast pool may constitute a first line of response to tissue damage, rapidly differentiating to generate new ECs, while giving time to stem cells to progress through the cell cycle.

Our culture system can also be used in combination with temperature-sensitive gene induction or knock-down tools, thus expanding its applications. When genetically stimulated by the expression of constitutively active *Ras*^*G12V*^, ISCs proliferated rapidly (Figure 4 and Video 7). Similarly, co-overexpression of *stg* and *CycE* also resulted in ISC proliferation. When reconstructing cell lineages, we observed both symmetric and asymmetric cell divisions. We defined the former as a division event giving rise to two daughter cells which were then both seeing dividing further. Asymmetric divisions where instead functionally defined as giving rise to two daughter cells, only one of which could be seen dividing further, while the other differentiated or remained quiescent for the remainder of the imaging session, which had to last at least 6h after the sister’s mitotic event. By analyzing the reconstructed lineages, we found that upon *Ras*^*G12V*^ stimulation most lineages did not present symmetric divisions. Interestingly, this is accordance to the previously described prevalence of asymmetric division events in normal intestines^47,66^. This suggests that the EGFR-Ras-Erk pathway may have a role in differentiation. Indeed, several progenitor cells, when stimulated by *Ras*^*G12V*^, did not divide, but rapidly grew in nuclear size and lost *esg* expression, which is indicative of EB to EC differentiation and a similar phenotype to what previously reported^57^. However, as cells were continuously stimulated by *Ras*^*G12V*^, it is unclear whether the Ras pathway had a direct effect on generating asymmetry during mitotis, or simply stimulated the differentiation of daughter cells.

In analyzing ISC lineages from live analysis we defined 8 ISC division events as symmetric and 17 as asymmetric. Reconstructing 3D profiles, we found that in most cases, right after a mitotic event both sister cells laid flat against the basal layer of visceral muscle, with no difference between symmetric and asymmetric mitotic events. However, it must be pointed out that to limit phototoxicity we imaged midguts with a time interval of 1h, a much longer interval than what previously used for spindle analyses^21,22^. Given that mitoses were measured to last between 30 to 60 minutes^21^, this means we only rarely managed to image the actual mitotic event, but instead captured the result of it. It is therefore possible that mitotic spindles were oriented differently in asymmetric and symmetric divisions, but that this difference in orientation was lost after cytokinesis.

One major difference that we did observe between symmetric and asymmetric ISC divisions was in the behavior of sister cells. Daughters of asymmetric divisions remained close to one another for several hours after mitosis. This is significant as it is known that cell-cell contacts between progenitor cells are required for promoting differentiation^1,2^. Indeed, interactions between Notch on the surface of the EB and Delta expressed on the ISC surface is a strong promoter of EB to EC differentiation^1,2,51^. It was previously shown that EB differentiation via N activation requires several hours to resolve^21^. This time frame matches our observations, which show asymmetric sister pairs remaining in contact for at least ~3-5 hours after division. Adherens junctions may be affecting these cell contacts as strong levels of shotgun (e-cadherin) and armadillo (β-catenin) are found in between ISC and EB pairs^65^. As EGFR signaling is known to impact adherens junctions remodeling^53,70,71^, this could help explain the prevalence of asymmetric mitosis in ISC lineages stimulated by *Ras*^*G12V*^ expression. Symmetric pairs that generated two proliferative cells had the opposite behavior, and moved apart right after division. This is also significant since it would result in the dispersion of stem cells through the midgut. As symmetric divisions are known to occur during adaptive growth of the intestine, especially in the days after eclosion, this behavior could help explain how ISCs space themselves uniformly in the epithelium^66^.

Despite these successful applications, the system we developed still has limitations. The optimized dissection procedure limits midgut damage, but does not completely eliminate the risk. The agarose pads, albeit thin, can interfere with high powered objectives with short working distances (*e*.*g*. 40X and above). Using thinner agarose pads could help to reduce the required working distance, but may result in hypoxia. This in turn could be solved by increasing the oxygen concentration using microscope incubation chambers equipped with an atmosphere control unit. Midgut survival *ex vivo* also seems to be limited by the growth of enteric bacteria, especially since, once dissected, midguts cannot properly move food through the intestine and defecate. Indeed, we observed that the visceral muscle, which is not in direct contact with luminal contents, could survive and contract regularly for up to a week in optimal conditions, despite the death of the adjacent epithelium. Generating axenic flies may help to further extend the survival of the midgut epithelium *ex vivo*. Newer, gentler imaging technologies such as light-sheet microscopy, could also improve survival during live-imaging sessions by reducing phototoxicity. Lastly, it’s also possible that different organs may have specific requirements in terms of media composition and additives, in which case tailoring culture media to a specific organ may be beneficial. Indeed, even the same organ can have different requirements in male and female flies^72–74^.

Nonetheless, the increased survival *ex vivo* our protocol allows is significant and we believe it can enable experiments that will lead to a better understanding of the mechanisms that mediate epithelial homeostasis, such as the regulation of asymmetric and symmetric ISC division events. Our protocol may also provide a platform to dissect inter-organ interactions, given the positive effect that co-culture with ovaries and fat bodies had on midgut survival (Figure 2A). Moreover, Malpighian tubules, hearts, and ovaries did show increased survival when cultured with our protocol (Figure 8 and Videos 17-20). Finally, we believe that the possibility to visualize the effects of gene induction or silencing in real-time using fluorescent markers will be very useful for dissecting the roles of specific signaling pathway components and in modeling human disease.

## MATERIALS AND METHODS

### *Drosophila* stocks

*w*^*1118*^ (Bloomington Drosophila Stock Center 3605)

*esg-Gal4 tubGal80ts UAS-GFP / CyO; UAS-flp Act>CD2>Gal4 / TM6B* (PMID: 19563763)

*esg-Gal4 Tub-Gal80*^*TS*^ *UAS-nlsGFP / CyO*

*mex-Gal4 Tub-Gal80*^*TS*^ *UAS-GFP / CyO*

*esg-Gal4 UAS-GFP*

*esg-Gal4 Tub-Gal80*^*TS*^ *UAS-nlsGFP / CyO*; *His2Av-mRFP / TM6B*

*His2Av-mRFP* (Bloomington Drosophila Stock Center 23650)

*UAS-N*^*RNAi*^ */ TM6B* (PMID: 26237646)

*UAS-Ras*^*G12V*^ */ CyO* (PMID: 21167805)

*UAS-stg, UAS-CycE* (PMID: 24975577)

*PCNA-GFP* (Stefano Di Talia, Duke University Medical Center, USA)

*His2Av-mRFP; PCNA-GFP*

### Fly rearing

Flies were raised on standard cornmeal and molasses fly food. Prior to dissection, flies were flipped to fresh vials without live yeast daily for 3 days at 18°C to reduce the load of commensal bacteria. On the morning of the dissection, flies were fed a sucrose 0.05% aqueous solution on a cotton pad at room temperature (25°C) for 4-6h to clear most luminal contents. This also helped to reduce accumulation of food in the posterior section of the gut, which could lead to the mechanical stress of the epithelium. For SDS feeding experiments, flies were fed overnight either standard food or food mixed with SDS to a final concentration of 0.2% v/w. Both foods were also mixed with a blue food-safe dye to control for feeding. The following morning, flies were fed a sucrose 0.05% aqueous solution on a cotton pad to clear most of the luminal SDS.

### Modified Schneider’s medium for adult *Drosophila* tissues

Schneider’s medium (Genesee Scientific, 25-515) was modified by adding the following reagents to the stated final concentrations: 1mM trisodium citrate dihydrate (ThermoFisher Scientific, BP327), 91.2mM sodium chloride (Sigma-Aldrich, S9888), 55.8mM D-trehalose anhydrous (Sigma-Aldrich, T0167), 10mM glutamine (Gibco, 25030), and 2mM N-acetyl cysteine (Sigma-Aldrich, A7250) (Table 1). Glutamine needs to be added only if the batch of Schneider’s medium used is glutamine-free. Medium was then filtered using 0.22µm syringe filters (VWR, 28145) and stored at 4°C. See Supplementary File 1 for recipe. This medium was used without additives during dissection, agarose gel stock preparation, and as a base for fly extract.

For fly extract, well-fed female flies were anesthetized on ice. Using mortar and pestle, flies were homogenized on ice in the presence of 10µl per mg of flies of modified Schneider’s medium (as described above) with bovine serum albumin (BSA) added to 1%. The homogenate was centrifuged at 0.6G and 4°C for 10’. Supernatant was saved and fly carcasses discarded. The centrifugation step was repeated 3 times until all solid fly residues were eliminated. Extract was heat inactivated by heating at 60°C for 5’, then centrifuged at 0.6G and 4°C for 10’. Supernatant was saved and filtered using 0.22µm syringe filters. Extract was aliquoted and stored at −20°C before use.

For live-imaging, 100% fly extract prepared in modified Schneider’s medium was used as a base for the complete culture medium. Fly extract was slowly thawed at 4°C, then 10% fetal calf serum (Gibco, 26140079), 1:100 Antibiotic-Antimycotic (ThermoFisher Scientific, 152400062), 100µg/ml Ampicillin (Fisher Scientific, AC611770250), and 25µg/ml Chloramphenicol (Fischer Scientific, BP904-100) were added. To suppress peristaltic movements, 10µg/ml Isradipine (SigmaAldrich, I6658) was added to the complete medium immediately before imaging.

### Sample preparation for long-term culture and live imaging

1. Prior to dissection, a 35mm dish with lockable lid (ibidi®, µ-Dish 35 mm low, 80136) is prepared by first placing a thin wet paper tissue around its inner rim to reduce evaporation during long imaging sessions (Figure 1O, left panel);
2. Agarose pads are then cast by spreading 2µl of low gelling temperature agarose (Sigma Aldrich, A9414), heated to 70°C, over four 4mm areas in the observation region of the dish. For each dish, 4 pads can be easily cast (Figure 1O, left panel). The agarose solution is prepared from powder as a 1% stock in modified Schneider’s medium without additives and stored at 4°C in 200µl aliquots. Aliquots can be melted and re-gelled several times, provided evaporation is not excessive;
3. Dishes are then stored at room temperature while midguts are isolated by dissection;
4. A small amount of medium is then added to the top of each agarose pad to facilitate the transfer of midguts. These are transferred very carefully, by holding them in a drop of liquid in between the grasping ends of a forceps. The drop is then touched to the top of an agarose pad, gently depositing the midgut trapped in it. Care must be exercised to ensure that the midgut rests entirely within the drop of liquid, without touching the dry surfaces of the forceps, to which it may stick. Each agarose pad can house up to 3 guts;
5. Once all midguts have been transferred, liquid from the top of the agarose pads is removed as much as possible using forceps, while leaving a small amount to avoid desiccation of the intestines. Midguts are then gently repositioned for proper imaging, if required;
6. Guts are then covered with a thin layer of low gelling temperature agarose 0.5% cooled to 37°C (Figure 1O, middle panel). The layer must be just enough to cover the midguts’ surface (~1µl per pad);
7. The sample is incubated for 5 min at room temperature before the agarose structures are connected between them and to the sides of the observation area by creating agarose bridges with 0.5% low gelling agarose (Figure 1O, middle panel). This increases the stability of the overall sample, facilitating its transport in case the sample has to be prepared at some distance from the microscope that will be used to image it. If required, agarose domes can also be strengthened with an additional thin layer of agarose;
8. After 10 min, the agarose will have solidified and 120µl of complete culture medium can be carefully added to the sample (Figure 1O, right panel). The small volume is enough so that all midguts will receive nutrients throughout the culture duration, while ensuring that the uppermost surface of the agarose structures is not submerged by liquid, creating a liquid-air interface;
9. Finally, ovaries and fly abdomens that were dissected along with the midguts are added to the culture, free-floating in between the agarose pads. The sample will thereby be ready for imaging;

### Optimized dissection to avoid damaging of midguts

1. To reduce the risk of contamination of the culture by bacteria residing on the animal exterior, CO_2_ anesthetized flies are surface sterilized by submerging them in 70% ethanol for 2 min and then in 50% bleach for 1 min. Most flies survive this treatment;
2. Flies are then washed and stored in 1X PBS modified as they are dissected one by one in modified Schneider’s medium without additives. Since this step is fast (< 2 min), using complete medium based on fly extract is not required;
3. Using micro-scissors the head is removed with a clean cut, thus ensuring that the crop and proventriculus still reside in the fly thorax (Figure 1B,C; Video 1);
4. The cuticle around the anus is pulled, exposing the hindgut (Figure 1D);
5. Holding the fly gently with forceps around the thorax-abdomen junction, the soft ventral abdominal cuticle is ripped using another forceps, pulling it along the length of the fly towards the anus, thus exposing the midgut (Figure 1E,F; Video 1);
6. The abdomen is then gently separated from the thorax (Fig. 1G; Video 1);
7. The crop is gently pinched and pulled out of the thorax, thus freeing the anterior midgut along with it (Figure 1H,K yellow arrowhead; Video 1);
8. The midgut is gently freed from the abdominal cuticle (Figure 1J; Video 1). Care has to be exercised at this step as many trachea filaments connect the midgut to ovaries and abdominal walls;
9. Crop and malpighian tubules are cut away using micro-scissors, and the hindgut is similarly removed just below its connection to the midgut (Figure 1K-M; Video 1). This step is necessary as both Malpighian tubules and the ampulla connected to the hindgut are quite sticky and make transferring the guts to the imaging dish difficult. If desired, however, Malpighian tubules can be transferred to agarose pads for imaging, either detached or still connected to midguts;
10. Once dissected, a midgut can then be transferred to a well containing modified Schneider’s medium with 10µg/ml Isradipine. Ovaries and the fly abdomen are transferred to this well as well;
11. Once all midguts have been dissected, they can be carefully transferred to the agarose pads for sample preparation. To avoid damage during transfer, it is recommended that midguts be moved by keeping them in a small drop of liquid in between the prongs of a forceps;
12. For localized damage experiments, midguts were poked in their posterior section using an electrolytically sharpened tungsten needle once transferred to the agarose pads. To keep guts still during the procedure, the hidgut section can be clamped with forceps held in the non-dominant hand. For this, a longer hidgut section can be preserved (see point 9). It is imperative to avoid perforating the peritrophic matrix, thus preventing the contamination of the culture from commensal bacteria. For this, guts are not perforated perpendicularly to the epithelium, but at a ~45° angle. Alternatively, guts can be carefully scratched repeatedly in the same spot to produce a tear;

### Long-term live-imaging of explanted adult *Drosophila* midguts

For imaging of midgut, hearts, and Malpighian tubules we used an inverted widefield Nikon Ti Eclipse microscope equipped with an incubation chamber (Okolab) for moisture and temperature control, a CoolSnap HQ2 camera (Photometrics), a SOLA LED light engine (Lumencore), and a Prior motorized stage. To limit phototoxicity, albeit at the cost of reduced signal/noise ratio, we used a 4x neutral density filter, limited light intensity to 5%, and limited exposure times to 100-150ms and 200-300ms for GFP and RFP signals, respectively. We also tested our culture conditions using a Leica SP8 confocal microscope also equipped with an incubation chamber (Okolab) for moisture and temperature control. While laser power and dwell time were minimized to reduce phototoxicity, we found this setup less viable than the widefield microscope. Nonetheless, the better resolution in the Z axis proved beneficial for the imaging of follicles. For both setups, videos were captured at room temperature (25°C) or at 29°C for temperature-sensitive gene induction experiments using a 20X air objective (APO 20X NA 0.75 WD 1). Multipoint acquisition was used to image the R4-5 posterior midgut section of 12 intestines during each imaging section. For each midgut, a 45µm Z-stack was captured using a 3µm Z-step. Focus was manually checked and corrected during the course of the imaging session with the Nikon Ti Eclipse, or using a contrast-based autofocus routine for the Leica SP8. Frame rate was typically one full multi-channel Z-stack/midgut/hour. For the imaging of Malpighian tubules and ovaries contractions and beating hearts, the frame rate was 1 slice every 0.08ms instead.

### NucGreen analysis

For cell survival experiments, guts from W^1118^ flies were cultured in media containing NucGreen (ThermoFisher Scientific, R37109) diluted 1:20. Midguts were imaged once a day for 3 days, using a 10X air objective (Plan Fluor 10X NA 0.3 WD 16) to capture a stitched z-stack of the whole organ with z-steps of 20µm. Images were normalized to their median pixel value to account for changes in background, then midgut areas were manually selected from maximum intensity projections. The NucGreen signal was computed as the sum of pixel values in the selected areas, normalized to the first time-point in the series.

### GFP level quantification

Midguts were explanted from 5-15 days old flies reared at 18°C and cultured at 29°C for 24h. Midguts were then carefully removed from the culture setup and fixed in para-formaldehyde 6% in PBS for 30 minutes at room temperature. Similarly, *in vivo* control flies were shifted to 29°C for 24h, then dissected and their midguts fixed. Fixed intestines were stained for DAPI 0.1mg/ml (Sigma-Aldrich, 10236276001) in PBS with 0.1% Triton-X100, then mounted with VECTASHIELD® antifade mounting medium (Vector Laboratories, H-1000-10). The posterior region of the intestines was imaged and the GFP and DAPI signals were thresholded using Otsu’s method^75^. Regions of overlap between the two tresholded channels were used as a mask to calculate the mean GFP expression of each cell of interest in the imaged field. For each intestine, the values of each measured cell were averaged to express a mean fluorescence for the whole intestine.

For GFP induction tests, malpighian tubules were detached from midguts using micro-scissors and cultured in sandwiched agarose structures as described above. Tubules were cultured at 18°C for 24h then at 29°C. Images were captured daily. Individual GFP^+^ cells from 48h images were identified in previous time-points and their GFP intensity measured.

### Cell tracking and analysis

Image analysis was performed using either ImageJ or Python (v3.7.10). Each time-lapse movie was divided at the first time point into overlapping regions of interest that were then used for automatic local registration. This helps account for local movements and deformations of the intestinal epithelium which complicate the following image analysis step. Registration was based on cross-correlation of a region of interest with the next frame of the time-series. Individual ISC and their progeny were then manually tracked using a custom ImageJ macro, selecting each cell by drawing their outline at their most in-focus z slice. By identifying all cells between successive time-points, we avoided lineage assignment errors even with long time-intervals between frames.

Cells were then analyzed as follows:

- Lineages were reconstructed with a custom Python script using the cell positions in the imaged field across time annotated during manual tracking. When the former approach failed due to too complex rearrangements of cells within a lineage, lineages were manually annotated instead;
- Nuclear size was measured during manual tracking from the most in-focus z-slice using an upscaled image so as to achieve sub-pixel precision;
- Mean nuclear GFP intensity was measured during manual tracking and adjusted to the median value in a 50µm window around the cell of interest to account for changes in local background;
- Cell cycle duration was measured as the number of frames (each representing a 1h interval) from the appearance of a cell resulting from a mitotic event and its subsequent division;
- Cell motility was defined as the distance between the positions of a cell in two subsequent frames. A few errors during image registration resulted in improper measurements, which were then discarded. For this, outlier values were first identified using the interquartile range method, then confirmed on the registered movie file;
- For analysis of the angle between sister cells, the z-profile along the axis connecting the center of the two cells was reconstructed Python. Only cells in flat sections of epithelium in reference to the imaging plane (based on enterocytes’ nuclei positions) were considered. The pixel values of the stack along a line connecting the two cells were used to reconstruct the 3D profile. Once all profiles were reconstructed, the angle between the 3D centers of the sister cells was manually computed in reference to the imaging plane. Measurements were repeated three times and averaged.

## ACKNOWLEDGEMENTS

This work was supported by the Huntsman Cancer Foundation, and National Institutes of Health Grants R01 GM124434 and R35 140900 (to B.A.E.) and P30 CA042014. We thank the Bloomington Drosophila Stock Center for fly stocks. We thank the University of Utah Cell Imaging Core for its instruments and assistance.

## AUTHOR DETAILS

### Marco Marchetti

Department of Oncological Sciences, Huntsman Cancer Institute, University of Utah, Salt Lake City, United States

### Contribution

Conceptualization, Methodology, Software, Validation, Investigation, Resources, Data Curation, Writing – Original Draft Preparation, Writing – Review & Editing, Visualization.

### Competing interests

No competing interests declared.

### Chenge Zhang

Department of Oncological Sciences, Huntsman Cancer Institute, University of Utah, Salt Lake City, United States

### Contribution

Conceptualization, Validation, Investigation, Resources, Writing – Original Draft Preparation, Writing – Review & Editing, Visualization.

### Competing interests

No competing interests declared.

### Bruce Alexander Edgar

Department of Oncological Sciences, Huntsman Cancer Institute, University of Utah, Salt Lake City, United States

### Contribution

Conceptualization, Validation, Writing – Original Draft Preparation, Writing – Review & Editing, Visualization, Supervision, Project Administration, Funding Acquisition.

### Competing interests

No competing interests declared.

## COMPETING INTERESTS

The authors declare no competing interests.

## SUPPLEMENTARY DATA

**Supplementary File 1: Modified Schneider’s medium and stock solutions recipes**.

**Video 1: Dissection technique to minimize damage during midgut explantation**. See also Figure 1 and methods section.

**Video 2: 72h live-imaging of an undamaged intestine expressing a lineage tracing system under *esg***^***TS***^

**driver**. Maximum intensity projection. See also Figure 2C. Scale bar is 5µm.

**Video 3: 48h live-imaging of a healthy intestine expressing *His2Av*.*mRFP* (red) and *esg***^***TS***^**-driven GFP (Green) induced 24h prior to imaging**. Maximum intensity projection. See also Figure 3A. Scale bar is 20µm.

**Video 4: 48h live-imaging of an intestine from a SDS-fed fly expressing His2Av.mRFP (red) and *esg***^***TS***^**- driven GFP (Green) induced 24h prior to imaging. Maximum intensity projection**. See also Figure 3B. Scale bar is 20µm.

**Video 5: detailed view of Video 2 showing a progenitor cell (white outline) differentiating and replacing dying enterocytes (yellow circles) in an intestine from a SDS-fed fly expressing His2Av.mRFP (red) and *esg***^***TS***^**-driven GFP (Green) induced 24h prior to imaging**. Maximum intensity projection. See also Figure 3B’. Scale is 10µm.

**Video 6: 20h live-imaging of an intestine expressing *esg*-driven GFP (Green) damaged by needle poke 10’ prior to imaging**. Maximum intensity projection. See also Figure 3 – figure supplement 3B. Scale bar is 20µm.

**Video 7: 24h live-imaging of an undamaged intestine expressing His2Av.mRFP (red) and upd3-driven GFP (green)**. Maximum intensity projection. See also Figure 3 – figure supplement 1D.

**Video 8: 24h live-imaging of an intestine damaged using a tungsten needle and expressing His2Av.mRFP (red) and upd3-driven GFP (green**). Damaged area is marked by the yellow arrowhead. Maximum intensity projection. See also Figure 3 – figure supplement 1E.

**Video 9: 48h live-imaging of an undamaged intestine expressing His2Av.mRFP (red) and *esg***^***TS***^**-driven GFP (Green) and *N***^***RNAi***^ **induced 24h prior to imaging**. Maximum intensity projection. See also Figure Figure 3 – figure supplement 2A. Scale bar is 20µm.

**Video 10: 48h live-imaging of a damaged intestine expressing His2Av.mRFP (red) and *esg***^***TS***^**-driven GFP (Green) and *N***^***RNAi***^ **induced 24h prior to imaging**. Damaged area is marked by the yellow ellipses. Maximum intensity projection. See also Figure Figure 3 – figure supplement 2B. Scale bar is 20µm.

**Video 11: 48h live-imaging of a healthy intestine expressing His2Av.mRFP (red) and *esg***^***TS***^**-driven GFP (Green) induced after dissection**. Maximum intensity projection. See also Figure 4A. Scale bar is 20µm.

**Video 12: 48h live-imaging of an intestine expressing His2Av.mRFP (red) and *esg***^***TS***^**-driven GFP (Green) and *Ras***^***G12V***^ **induced after dissection**. Maximum intensity projection. See also Figure 4B. Scale bar is 20µm.

**Video 13: 48h live-imaging of an intestine expressing His2Av.mRFP (red) and *esg***^***TS***^**-driven GFP (Green), *stg*, and *CycE* induced after dissection**. Maximum intensity projection. See also Figure 4C. Scale bar is 20µm.

**Video 14: Example of ISC lineage. Arrowheads indicate cells in the lineage**. Maximum intensity projections of z-slices encompassing the cells in the lineage. See also Figure 5. Scale bar is 5µm.

**Video 15: Example of symmetric division. Arrowheads indicate cells in the lineage, with symmetric daughter pairs marked by green ones**. Maximum intensity projections of z-slices encompassing the cells in the lineage. See also Figure 6A. Scale bar is 5µm.

**Video 16: Example of asymmetric division**. Arrowheads indicate cells in the lineage, with asymmetric daughter pairs marked by red ones. Maximum intensity projections of z-slices encompassing the cells in the lineage. See also Figure 6C. Scale bar is 5µm.

**Video 17: Example of Malpighian tubule contracting after 3 days *ex vivo*. Note that the tubule is still connected to the midgut**. To avoid inhibiting contractions, Isradipine was not supplemented, resulting in the midgut rupturing near the imaged site, releasing visible debris. Scale bar is 50µm.

**Video 18: Example of heart lining the dorsal side of a dissected fly abdomen beating after 10 days ex vivo**. Scale bar is 50µm.

**Video 19: Example of ovary contracting after 3 days ex vivo**. Scale bar is 100µm.

**Video 20: Example of stage 4 follicle progressing to stage 5 *ex vivo* and starting to elongate and rotate along its long axis**. Maximum intensity projection of the 3 center-most z-slices. See also Figure8C. Scale bar is 10µm.

**Figure 2 – source data 1**. Raw data for Figure 2A, D, E. **Figure 3 – source data 1**. Raw data for Figure 3D, E. **Figure 4 – source data 1**. Raw data for Figure 4D, F. **Figure 6 – source data 1**. Raw data for Figure 6E-H. **Figure 7 – source data 1**. Raw data for Figure 7A, D, F.

**Figure 7 – figure supplement 1 – source data 1**. Raw data for Figure 7 – figure supplement 1A-F.

**Figure 8 – source data 1**. Raw data for Figure 8B.

**Source code 1**. This Python script is used for local registration of adult *Drosophlila* midgut fluorescent imaging time-lapses. Movies are converted to 8bit, normalized by subtracting mean and dividing by the standard deviation, converted to maximum intensity or focused projections, then divided in partially overlapping regions of interest (ROIs). Each ROI is then XY-registered via cross-correlation between frames and then the most in-focus Z slice is found. Finally, ROIs are exported.

Source code 2. ImageJ Macro designed for the manual tracking of cells in two-channel time-lapse images of adult *Drosophila* midguts. Images must be two-channel time-lapse z-stacks, with channels 1 and 2 being His2Av.mRFP and esg^TS^ > nlsGFP, respectively.

**Source code 3**. This Python script is used to parse files generated by the “Source code 2” ImageJ macro. First, the lineage is reconstructed based on relative cell distances between frames. For each cell in frame f, a new position in frame f+1 is assigned, based on the pool of annotated cell positions in f+1. All possible permutations between f and f+1 coordinates are computed and the the total distance between cell positions in frame f and their newly assigned positions in f+1 is calculated. The permutation for which this distance is minimum is then kept. If lineages are too complex (e.g. cells are moving and rearranging themselves), then the user can add two additional field to the input “.tsv” file (“CellID” and “MotherID” columns) indicating each cells ID (for each cell position) and its mother cell ID (can be stated once). The lineage data is then parsed and formatted in an easy to read format, then exported to a “.tsv” file.

**Figure 2 – Figure supplement 1:**
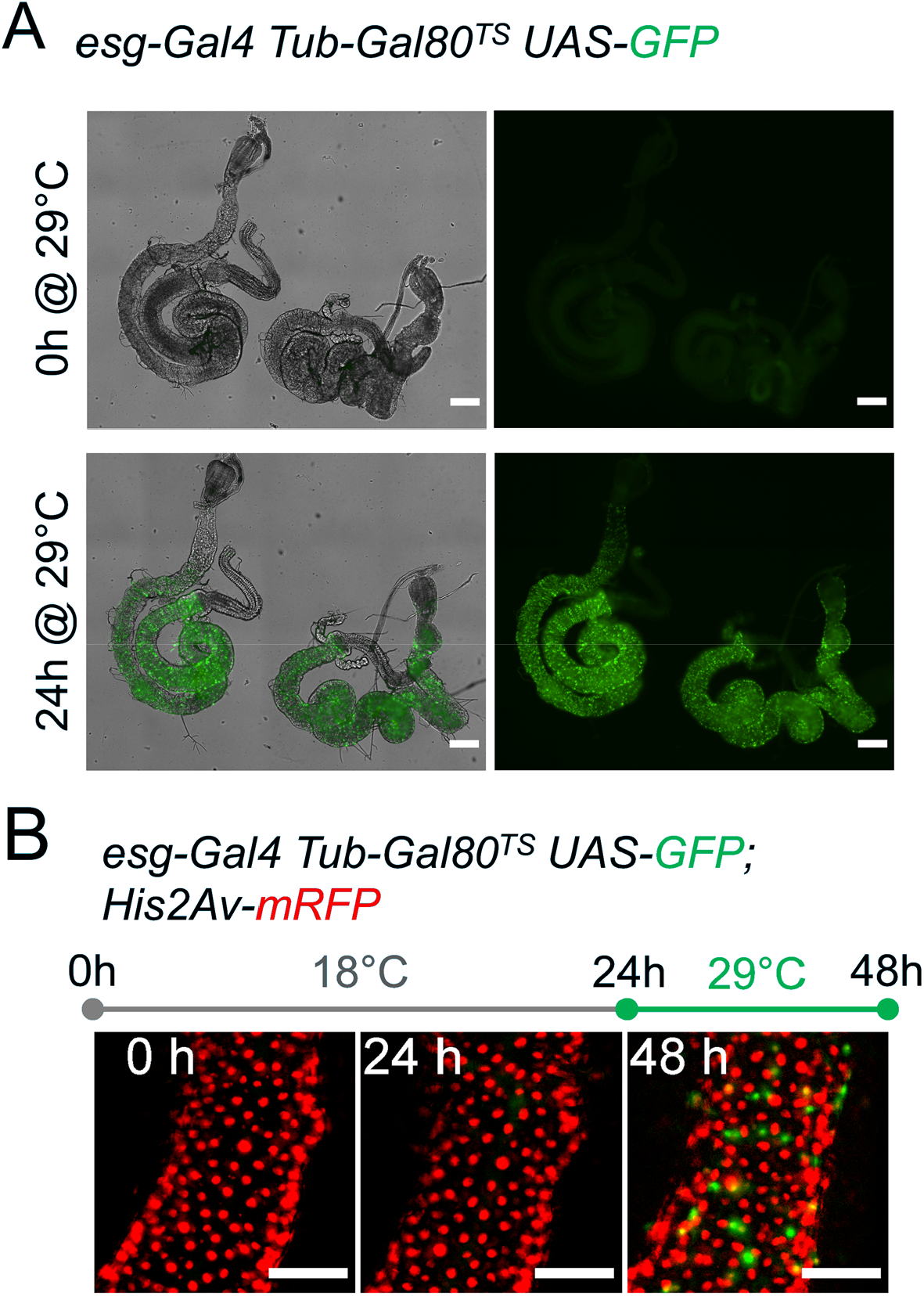
*Ex vivo* temperature sensitive gene expression of GFP. (A) Intestines explanted from flies raised at 18°C show no GFP expression. After explantation, guts were cultured at 29°C and started to express GFP driven by the *esg*^*TS*^ system. GFP channel is a maximum intensity projection. Scale bar is 200µm. (B) Intestines cultured at 18°C for 24h, then shifted to 29°C are still able to activate expression of GFP, showing that the epithelium remains healthy long-term. Images are maximum intensity projections. Scale bar is 50µm.

**Figure 3 – Figure Supplement 1:**
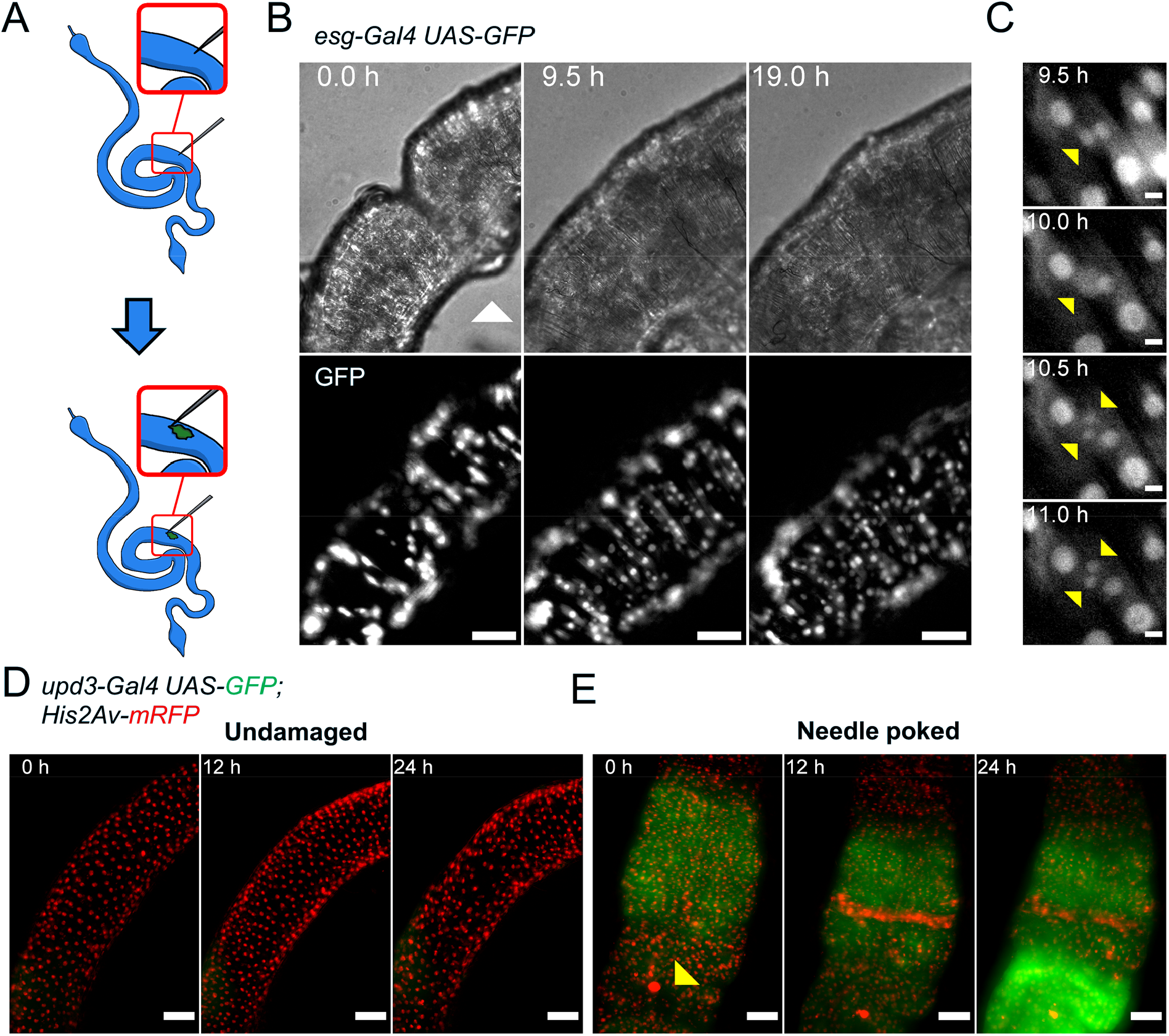
Tissue repair in midguts damaged *ex vivo*. (A) Diagram depicting the laceration of midgut tissues with a tungsten needle. (B) Time-lapse of an intestine expressing GFP via *esg*^*TS*^ driver and damaged with a tungsten needle (white arrowhead at time-point 0h). Note the gradual replacement of the epithelium by the proliferation/differentiation of GFP^+^ progenitor cells. Images are maximum intensity projections. See also Video 6. Scale bar is 50µm. (C) Example of stem cell dividing upon tissue damage from the intestine shown in panel B. Scale bar is 5µm. (D) Undamaged intestine expressing only low levels of upd3 (green). Images are maximum intensity projections. See also Video 7. Scale bar is 50µm. (E) Intestine damaged with the protocol described in panel (A) showing strong activation of the upd3 reporter (green) at the damaged area (marked by the yellow arrowhead). Images are maximum intensity projections. See also Video 8. Scale bar is 50µm.

**Figure 3 – Figure Supplement 2:**
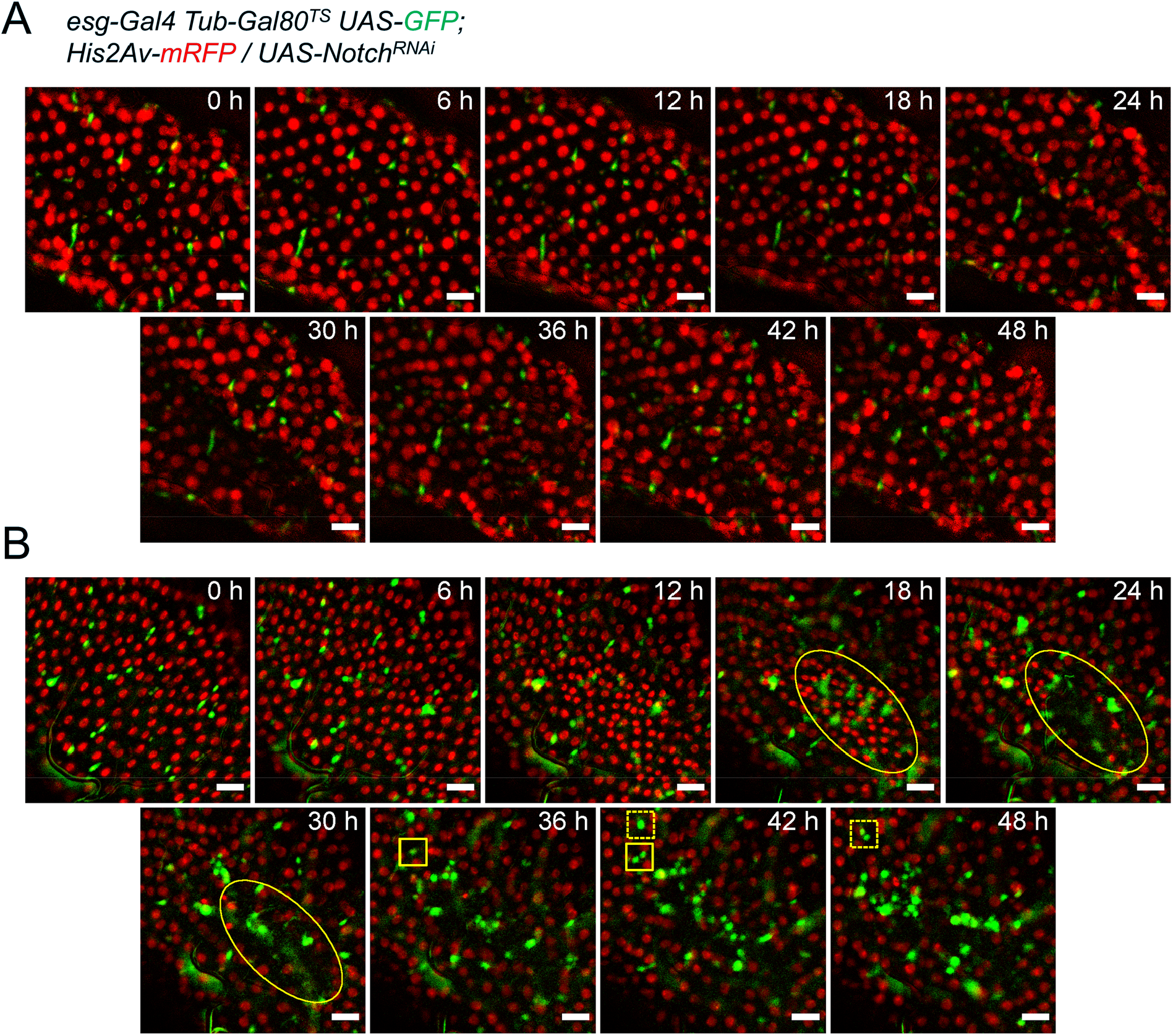
*Notch* knock-down induces tumorigenesis *ex vivo*. (A) Undamaged intestine expressing NRNAi under the *esg*^*TS*^ driver system showing no tumorigenesis. Images are maximum intensity projections. See also Video 9. Scale bar is 20µm. (B) Intestine expressing NRNAi under the *esg*^*TS*^ driver system accidentally damaged during dissection. The damaged area (yellow ellipses) shows massive cell death. In response, esg^+^ progenitor cells started to proliferate (see full and dashed yellow rectangles for examples, matching each between frames). Unlike in normal damage conditions, progenitor cells failed to differentiate (compare to Figure 3B). Images are maximum intensity projections. See also Video 10. Scale bar is 20µm.

**Figure 7 – Supplement 1.**
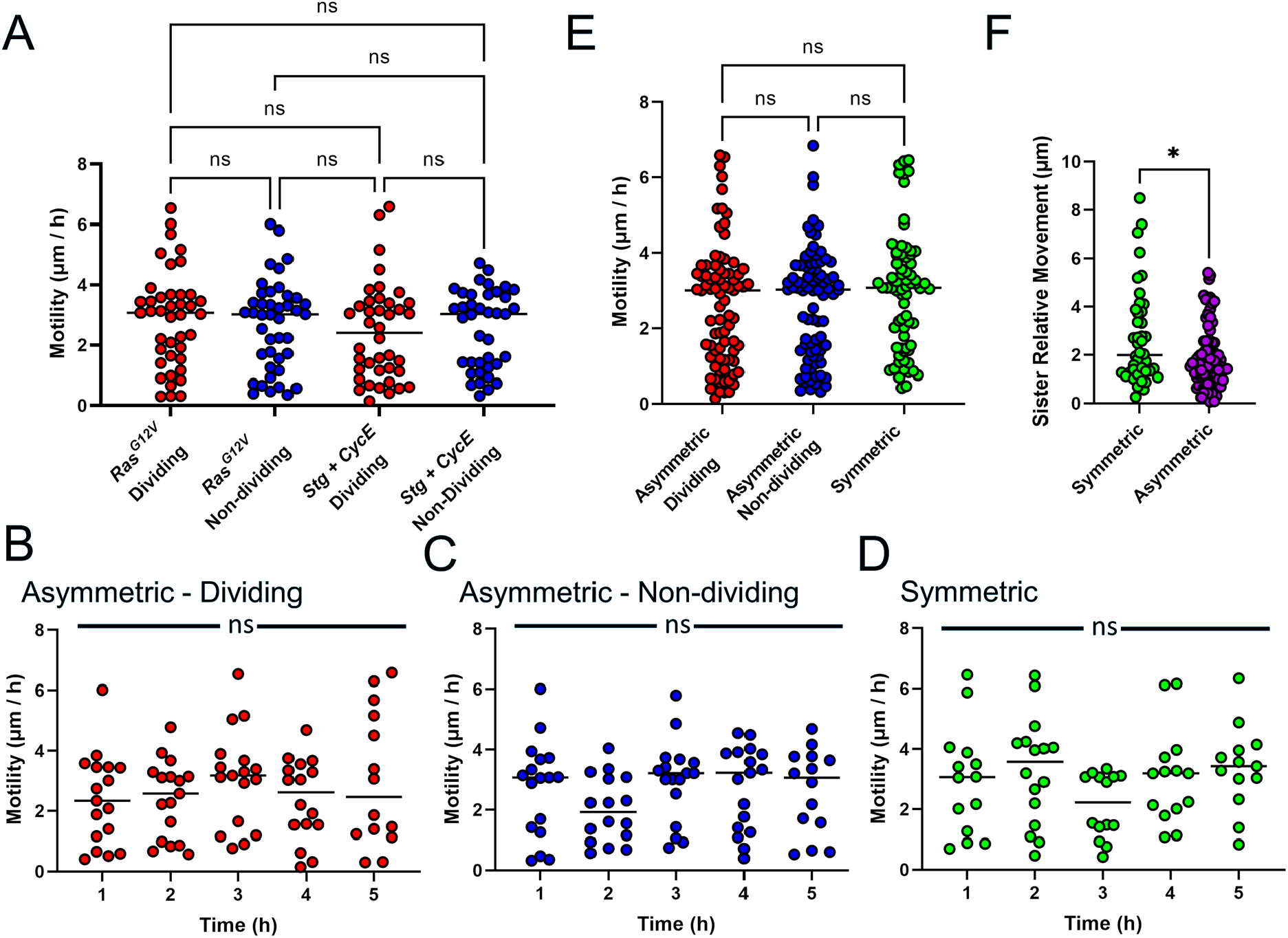
Lack of effect of genotype and time-point after mitosis on cell motility. For all graphs, red and blue denote data from dividing and non-dividing progenies of asymmetric mitoses, respectively, purple refers to data from all cells in asymmetric pairs, while green refers to data from the progeny of symmetric pairs. (A) The genotype used to promote ISC proliferation does not result in differences in cell motility in the dividing (red) or non-dividing (blue) progeny of asymmetric mitoses (Two-way Anova and Tukey multiple comparisons test). (B-D) Cell motility calculated as the distance travelled by a cell between two subsequent time-points. X-axis indicates the time passed from the first appearance of a cell. Cell motility remains constant across time for the dividing (B), non-dividing (C) progeny of asymmetric divisions, as well as for the daughter cells generated by symmetric divisions (D) (Mixed-effects analysis was used instead of Anova to account for missing values due to cells proliferating and errors during image registration. Tukey’s test was used for multiple comparisons). (E) No difference in cell motility was found between the dividing (red) and non-dividing (blue) progeny of asymmetric mitoses and the daughter cells generated by symmetric divisions (green). (F) Movements of cells relative to their sisters for symmetric and asymmetric pais. Only the first 6 time-points after the first appearance of each pair was used for this analysis (Mann-Whitney test). (ns, not significant, *, p < 0.05)

## Notes

### Competing Interest Statement

The authors have declared no competing interest.

## REFERENCES

1. Ohlstein, B. & Spradling, A. The adult Drosophila posterior midgut is maintained by pluripotent stem cells. Nature 439, 470–474 (2006).

2. Micchelli, C. A. & Perrimon, N. Evidence that stem cells reside in the adult Drosophila midgut epithelium. Nature 439, 475–479 (2006).

3. Jiang, H. et al. Cytokine/Jak/Stat signaling mediates regeneration and homeostasis in the Drosophila midgut. Cell 137, 1343–1355 (2009).

4. Beebe, K., Lee, W.-C. & Micchelli, C. A. JAK/STAT signaling coordinates stem cell proliferation and multilineage differentiation in the Drosophila intestinal stem cell lineage. Dev. Biol. 338, 28–37 (2010).

5. Patel, P. H. et al. Damage sensing by a Nox-Ask1-MKK3-p38 signaling pathway mediates regeneration in the adult Drosophila midgut. Nat. Commun. 10, 1–14 (2019).

6. Sato, T. et al. Single Lgr5 stem cells build crypt-villus structures in vitro without a mesenchymal niche. Nature 459, 262–265 (2009).

7. Robb, J. A. MAINTENANCE OF IMAGINAL DISCS OF DROSOPHILA MELANOGASTER IN CHEMICALLY DEFINED MEDIA. J. Cell Biol. 41, 876–885 (1969).

8. Zartman, J., Restrepo, S. & Basler, K. A high-throughput template for optimizing Drosophila organ culture with response-surface methods. Development 140, 667–674 (2013).

9. Handke, B., Szabad, J., Lidsky, P. V., Hafen, E. & Lehner, C. F. Towards Long Term Cultivation of Drosophila Wing Imaginal Discs In Vitro. PLOS ONE 9, e107333 (2014).

10. Tsao, C.-K., Ku, H.-Y., Lee, Y.-M., Huang, Y.-F. & Sun, Y. H. Long Term Ex Vivo Culture and Live Imaging of Drosophila Larval Imaginal Discs. PLOS ONE 11, e0163744 (2016).

11. Strassburger, K. et al. Oxygenation and adenosine deaminase support growth and proliferation of ex vivo cultured Drosophila wing imaginal discs. Development 144, 2529–2538 (2017).

12. Siller, K. H., Serr, M., Steward, R., Hays, T. S. & Doe, C. Q. Live Imaging of Drosophila Brain Neuroblasts Reveals a Role for Lis1/Dynactin in Spindle Assembly and Mitotic Checkpoint Control. Mol. Biol. Cell 16, 5127–5140 (2005).

13. Rabinovich, D., Mayseless, O. & Schuldiner, O. Long term ex vivo culturing of Drosophila brain as a method to live image pupal brains: insights into the cellular mechanisms of neuronal remodeling. Front. Cell. Neurosci. 0, (2015).

14. Fichelson, P. et al. Live-imaging of single stem cells within their niche reveals that a U3snoRNP component segregates asymmetrically and is required for self-renewal in Drosophila. Nat. Cell Biol. 11, 685–693 (2009).

15. Morris, L. X. & Spradling, A. C. Long-term live imaging provides new insight into stem cell regulation and germline-soma coordination in the Drosophila ovary. Dev. Camb. Engl. 138, 2207–2215 (2011).

16. Reilein, A., Cimetta, E., Tandon, N. M., Kalderon, D. & Vunjak-Novakovic, G. Live imaging of stem cells in the germarium of the Drosophila ovary using a reusable gas-permeable imaging chamber. Nat. Protoc. 13, 2601–2614 (2018).

17. Cheng, J. & Hunt, A. J. Time-lapse Live Imaging of Stem Cells in Drosophila Testis. Curr. Protoc. Stem Cell Biol. CHAPTER, Unit-2E.2 (2009).

18. Deng, H., Gerencser, A. A. & Jasper, H. Signal integration by Ca2+ regulates intestinal stem cell activity. Nature 528, 212–217 (2015).

19. Xu, C., Luo, J., He, L., Montell, C. & Perrimon, N. Oxidative stress induces stem cell proliferation via TRPA1/RyR-mediated Ca2+ signaling in the Drosophila midgut. eLife 6, e22441 (2017).

20. He, L., Si, G., Huang, J., Samuel, A. D. T. & Perrimon, N. Mechanical regulation of stem cell differentiation through stretch-activated Piezo channel. Nature 555, 103–106 (2018).

21. Martin, J. L. et al. Long-term live imaging of the Drosophila adult midgut reveals real-time dynamics of division, differentiation and loss. eLife 7, e36248 (2018).

22. Hu, D. J.-K. & Jasper, H. Control of Intestinal Cell Fate by Dynamic Mitotic Spindle Repositioning Influences Epithelial Homeostasis and Longevity. Cell Rep. 28, 2807-2823.e5 (2019).

23. Pasco, M. Y. & Léopold, P. High Sugar-Induced Insulin Resistance in Drosophila Relies on the Lipocalin Neural Lazarillo. PLoS ONE 7, e36583 (2012).

24. Dus, M., Ai, M. & Suh, G. S. B. Taste-independent nutrient selection is mediated by a brain-specific Na+/solute co-transporter in Drosophila. Nat. Neurosci. 16, 526–528 (2013).

25. Park, S. et al. A Genetic Strategy to Measure Circulating Drosophila Insulin Reveals Genes Regulating Insulin Production and Secretion. PLoS Genet. 10, e1004555 (2014).

26. Tennessen, J. M., Barry, W., Cox, J. & Thummel, C. S. Methods for studying metabolism in Drosophila. Methods San Diego Calif 68, 105–115 (2014).

27. Matsushita, R. & Nishimura, T. Trehalose metabolism confers developmental robustness and stability in Drosophila by regulating glucose homeostasis. Commun. Biol. 3, 1–12 (2020).

28. Singleton, K. & Woodruff, R. I. The Osmolarity of Adult Drosophila Hemolymph and Its Effect on Oocyte-Nurse Cell Electrical Polarity. Dev. Biol. 161, 154–167 (1994).

29. Naikkhwah, W. & O’Donnell, M. J. Salt stress alters fluid and ion transport by Malpighian tubules of Drosophila melanogaster: evidence for phenotypic plasticity. J. Exp. Biol. 214, 3443–3454 (2011).

30. MacMillan, H. A. et al. Parallel ionoregulatory adjustments underlie phenotypic plasticity and evolution of Drosophila cold tolerance. J. Exp. Biol. 218, 423–432 (2015).

31. MacMillan, H. A., Andersen, J. L., Davies, S. A. & Overgaard, J. The capacity to maintain ion and water homeostasis underlies interspecific variation in Drosophila cold tolerance. Sci. Rep. 5, 1–11 (2015).

32. Olsson, T. et al. Hemolymph metabolites and osmolality are tightly linked to cold tolerance of Drosophila species: a comparative study. J. Exp. Biol. 219, 2504–2513 (2016).

33. Schneider, I. DIFFERENTIATION OF LARVAL DROSOPHILA EYE-ANTENNAL DISCS IN VITRO. J. Exp. Zool. 156, 91–103 (1964).

34. Schneider, I. Histology of larval eye-antennal disks and cephalic ganglia of Drosophila cultured in vitro. J. Embryol. Exp. Morphol. 15, 271–279 (1966).

35. Schneider, I. Cell lines derived from late embryonic stages of Drosophila melanogaster. J. Embryol. Exp. Morphol. 27, 353–365 (1972).

36. Schneider, I. & Blumenthal, A. B. Drosophila cell and tissue culture. in The Genetics and Biology of Drosophila. 265–315 (1978).

37. Davis, K. T. & Shearn, A. In Vitro Growth of Imaginal Disks from Drosophila melanogaster. Science 196, 438–440 (1977).

38. Britton, J. S. & Edgar, B. A. Environmental control of the cell cycle in Drosophila: nutrition activates mitotic and endoreplicative cells by distinct mechanisms. Dev. Camb. Engl. 125, 2149–2158 (1998).

39. Brand, A. H. & Perrimon, N. Targeted gene expression as a means of altering cell fates and generating dominant phenotypes. Dev. Camb. Engl. 118, 401–415 (1993).

40. McGuire, S. E. Spatiotemporal Rescue of Memory Dysfunction in Drosophila. Science 302, 1765–1768 (2003).

41. Liang, J., Balachandra, S., Ngo, S. & O’Brien, L. E. Feedback regulation of steady-state epithelial turnover and organ size. Nature 548, 588–591 (2017).

42. Li, Q. et al. Ingestion of Food Particles Regulates the Mechanosensing Misshapen-Yorkie Pathway in Drosophila Intestinal Growth. Dev. Cell 45, 433-449.e6 (2018).

43. Amcheslavsky, A., Jiang, J. & Ip, Y. T. Tissue Damage-Induced Intestinal Stem Cell Division in Drosophila. Cell Stem Cell 4, 49–61 (2009).

44. Patel, P. H., Dutta, D. & Edgar, B. A. Niche Appropriation by Drosophila Intestinal Stem Cell Tumors. Nat. Cell Biol. 17, 1182–1192 (2015).

45. Buchon, N., Broderick, N. A., Poidevin, M., Pradervand, S. & Lemaitre, B. Drosophila intestinal response to bacterial infection: activation of host defense and stem cell proliferation. Cell Host Microbe 5, 200–211 (2009).

46. Reiff, T. et al. Notch and EGFR regulate apoptosis in progenitor cells to ensure gut homeostasis in Drosophila. EMBO J. 38, e101346 (2019).

47. de Navascués, J. et al. Drosophila midgut homeostasis involves neutral competition between symmetrically dividing intestinal stem cells. EMBO J. 31, 2473–2485 (2012).

48. Lin, G., Xu, N. & Xi, R. Paracrine Wingless signalling controls self-renewal of Drosophila intestinal stem cells. Nature 455, 1119–1123 (2008).

49. Shaw, R. L. et al. The Hippo pathway regulates intestinal stem cell proliferation during Drosophila adult midgut regeneration. Dev. Camb. Engl. 137, 4147–4158 (2010).

50. Strand, M. & Micchelli, C. A. Quiescent gastric stem cells maintain the adult Drosophila stomach. Proc. Natl. Acad. Sci. 108, 17696–17701 (2011).

51. Ohlstein, B. & Spradling, A. Multipotent Drosophila Intestinal Stem Cells Specify Daughter Cell Fates by Differential Notch Signaling. Science 315, 988–992 (2007).

52. Apidianakis, Y., Pitsouli, C., Perrimon, N. & Rahme, L. Synergy between bacterial infection and genetic predisposition in intestinal dysplasia. Proc. Natl. Acad. Sci. U. S. A. 106, 20883–20888 (2009).

53. Buchon, N., Broderick, N. A., Kuraishi, T. & Lemaitre, B. Drosophila EGFR pathway coordinates stem cell proliferation and gut remodeling following infection. BMC Biol. 8, 152 (2010).

54. Biteau, B. & Jasper, H. EGF signaling regulates the proliferation of intestinal stem cells in Drosophila. Development 138, 1045–1055 (2011).

55. Jiang, H., Grenley, M. O., Bravo, M.-J., Blumhagen, R. Z. & Edgar, B. A. EGFR/Ras/MAPK signaling mediates adult midgut epithelial homeostasis and regeneration in Drosophila. Cell Stem Cell 8, 84–95 (2011).

56. Jin, Y. et al. EGFR/Ras Signaling Controls Drosophila Intestinal Stem Cell Proliferation via Capicua-Regulated Genes. PLoS Genet. 11, e1005634 (2015).

57. Xiang, J. et al. EGFR-dependent TOR-independent endocycles support Drosophila gut epithelial regeneration. Nat. Commun. 8, 15125 (2017).

58. Chen, J. et al. Transient Scute activation via a self-stimulatory loop directs enteroendocrine cell pair specification from self-renewing intestinal stem cells. Nat. Cell Biol. 20, 152–161 (2018).

59. Kohlmaier, A. et al. Src kinase function controls progenitor cell pools during regeneration and tumor onset in the Drosophila intestine. Oncogene 34, 2371–2384 (2015).

60. Goulas, S., Conder, R. & Knoblich, J. A. The Par complex and integrins direct asymmetric cell division in adult intestinal stem cells. Cell Stem Cell 11, 529–540 (2012).

61. Singh, S. R., Liu, W. & Hou, S. X. The Adult Drosophila Malpighian Tubules Are Maintained by Pluripotent Stem Cells. Cell Stem Cell 1, 191–203 (2007).

62. Wang, C. & Spradling, A. C. An abundant quiescent stem cell population in Drosophila Malpighian tubules protects principal cells from kidney stones. eLife 9, e54096 (2020).

63. Haigo, S. L. & Bilder, D. Global tissue revolutions in a morphogenetic movement controlling elongation. Science 331, 1071–1074 (2011).

64. Li, Z., Zhang, Y., Han, L., Shi, L. & Lin, X. Trachea-Derived Dpp Controls Adult Midgut Homeostasis in Drosophila. Dev. Cell 24, 133–143 (2013).

65. Choi, N. H., Lucchetta, E. & Ohlstein, B. Nonautonomous regulation of Drosophila midgut stem cell proliferation by the insulin-signaling pathway. Proc. Natl. Acad. Sci. 108, 18702–18707 (2011).

66. O’Brien, L. E., Soliman, S. S., Li, X. & Bilder, D. Altered Modes of Stem Cell Division Drive Adaptive Intestinal Growth. Cell 147, 603–614 (2011).

67. Jin, Y. et al. Intestinal Stem Cell Pool Regulation in Drosophila. Stem Cell Rep. 8, 1479–1487 (2017).

68. Dutta, D. et al. Regional Cell-Specific Transcriptome Mapping Reveals Regulatory Complexity in the Adult Drosophila Midgut. Cell Rep. 12, 346–358 (2015).

69. Hung, R.-J. et al. A cell atlas of the adult Drosophila midgut. Proc. Natl. Acad. Sci. U. S. A. 117, 1514–1523 (2020).

70. O’Keefe, D. D., Prober, D. A., Moyle, P. S., Rickoll, W. L. & Edgar, B. A. Egfr/Ras signaling regulates DE-cadherin/Shotgun localization to control vein morphogenesis in the Drosophila wing. Dev. Biol. 311, 25–39 (2007).

71. Robertson, F., Pinal, N., Fichelson, P. & Pichaud, F. Atonal and EGFR signalling orchestrate rok- and Drak-dependent adherens junction remodelling during ommatidia morphogenesis. Dev. Camb. Engl. 139, 3432–3441 (2012).

72. Hudry, B., Khadayate, S. & Miguel-Aliaga, I. The sexual identity of adult intestinal stem cells controls organ size and plasticity. Nature 530, 344–348 (2016).

73. Hudry, B. et al. Sex Differences in Intestinal Carbohydrate Metabolism Promote Food Intake and Sperm Maturation. Cell 178, 901-918.e16 (2019).

74. Ahmed, S. M. H. et al. Fitness trade-offs incurred by ovary-to-gut steroid signalling in Drosophila. Nature 584, 415–419 (2020).

75. Otsu, N. A Threshold Selection Method from Gray-Level Histograms. IEEE Trans. Syst. Man Cybern. 9, 62–66 (1979).

